# Non-phylogenetic identification of co-evolving genes for reconstructing the archaeal Tree of Life

**DOI:** 10.1101/2020.10.16.343293

**Authors:** L. Thibério Rangel, Shannon M. Soucy, João C. Setubal, Johann Peter Gogarten, Gregory P. Fournier

## Abstract

Assessing the phylogenetic compatibility between individual gene families is a crucial and often computationally demanding step in many phylogenomics analyses. Here we describe the Evolutionary Similarity Index (*I_ES_*) to assess shared evolution between gene families using a weighted Orthogonal Distance Regression applied to sequence distances. This approach allows for straightforward pairing of paralogs between co-evolving gene families without resorting to multiple tests, or *a priori* assumptions of molecular interactions between protein products from assessed genes. The utilization of pairwise distance matrices, while less informative than phylogenetic trees, circumvents error-prone comparisons between trees whose topologies are inherently uncertain. Analyses of simulated gene family evolution datasets showed that *I_ES_* was more accurate and less susceptible to noise than popular tree-based methods (Robinson-Foulds and geodesic distance) for assessing evolutionary signal compatibility, since it bypasses phylogenetic reconstruction and its inherent uncertainty. Applying *I_ES_* to a real dataset of 1,322 genes from 42 archaeal genomes identified eight major clusters of gene families with compatible evolutionary trends. Four of these clusters included genes with a taxonomic distribution across all archaeal phyla, while other clusters included a subset of taxa that do not map to generally accepted archaeal clades, indicating possible shared horizontal transfers by clustered gene families. We identify one strongly connected set of 62 genes from the same cluster, occurring as both single-copy and multiple homologs per genome, with compatible phylogenetic reconstructions closely matching previously published species trees for Archaea. An *I_ES_* implementation is available at https://github.com/lthiberiol/evolSimIndex.

## Introduction

Phylogenies reconstructed from single genes are known to poorly reflect the underlying history of whole genomes, as the detectable phylogenetic signal from an isolated locus cannot be extrapolated to represent whole genomes (Dagan and Martin 2006; Bapteste et al. 2009; Koonin et al. 2009). To ameliorate this effect, it has become common practice to estimate species’ evolutionary histories by concatenating multiple sequence alignments of genes from the core genome, i.e., single copy and widely conserved across sampled genomes, which greatly increases the number of sites available for phylogenetic inference. The preference towards using core genome sequences is due their expected resistance to horizontal gene transfer (HGT) (Thomas and Nielsen 2005; Sorek et al. 2007; Popa and Dagan 2011); however, despite the lower frequency of HGT among some gene families, it has been shown that horizontal exchange also affects genes from the core genome. In fact, the slow substitution rate and corresponding high sequence conservation of the core genome may even favor HGT, permitting increases in neutral and nearly-neutral HGT at the genus and species levels (Papke and Gogarten 2012; Shapiro et al. 2012).

Given this context, it is clear that more rigorous methods are needed to identify genes best reflecting the underlying vertical evolutionary signal in a group of species; such methods should seek to maximize the compatibility between evolutionary trends of the chosen gene families in order to provide a more robust basis for phylogenomic reconstruction. Many strategies have been proposed to assess similarities between the phylogenetic signals obtained by individual gene trees - e.g., Robinson-Foulds bipartition compatibility (RF) (Robinson and Foulds 1981) and geodesic distance (*D_geo_*) (Kimmel and Sethian 1998; Kupczok et al. 2008; Owen and Provan 2011) - as well as other methods that assess similarities between phylogenetic profiles (Pellegrini et al. 1999; Vert 2002; Barker and Pagel 2005; Liu et al. 2018). The majority of tree-based methods are based on straightforward comparisons between tree topologies (Kunin et al. 2005; Leigh et al. 2008; Puigbò et al. 2009; Mirarab et al. 2014; Gori et al. 2016). However, while an intuitive solution, comparisons between tree topologies require phylogenetic trees of all assessed gene families to be accurately reconstructed, adding a substantial computational cost to an already computationally demanding task. Furthermore, the vastness of tree space, combined with the inherent uncertainty of phylogenetic reconstruction, provides multiple sources of errors in tree-based evolutionary similarity assessments. Another method to assess the evolutionary compatibility of genes is based on similarities between patterns of presence and absence (phylogenetic profiles) of such genes among genomes of interest. While recent implementations displayed substantial improvements (Liu et al. 2018) when compared to the early ones, reliance on an initial reference tree hamper general applicability. Phylogenetic profile-based methods also do not assess divergencies between sequences of homologous genes, which limits the resolution of their results.

Accounting for uncertainty-based variations in tree topology (i.e., bipartition support) further increases the computational burden and decreases the resolution of the evaluated phylogenetic signal (e.g., collapsing low support bipartitions or weighing them based on support). A proposed solution to bypass the computational cost of tree similarity assessments is Pearson’s correlation coefficient (*r*) between evolutionary distance matrices (Goh et al. 2000; Pazos and Valencia 2001; Novichkov et al. 2004; Rangel et al. 2019). Unlike tree-based comparisons, methods based on Pearson’s *r* enable simple implementations to detect similar evolutionary signals between gene families with histories complicated by multiple homologs within genomes by estimating correlation coefficients using all possible pairings of paralogs between gene families (Gertz et al. 2003; Ramani and Marcotte 2003). Despite its application in protein-protein interaction studies, the sensitivity of Pearson’s *r* to noise in evolutionary distances and the granularity of its estimates have yet to be compared to those of tree-based metrics. Direct Coupling Analysis (DCA) has also been used to pair gene copies between possibly co-evolving gene families (Gueudré et al. 2016), but despite positive results the assumption that protein products of co-evolving genes must be directly interacting hampers its general applicability.

### New approaches

Given the shortcomings pointed out above, we propose the Evolutionary Similarity Index (*I_ES_*) as a metric for similarities between evolutionary histories based on weighted Orthogonal Distance Regression (wODR) between evolutionary pairwise distance matrices. We show that evolutionary similarity estimates from wODR display a linear correlation with performed stepwise perturbations simulated tree topologies. More common tree-based evolutionary similarity estimates, such as RF and *D_geo_*, tend to overestimate the impact of simulated topology changes, and consequently are significantly more susceptible to errors in evolutionary history reconstruction. As a case study of this new method, we assessed evolutionary similarities across 1,322 archaeal gene families and detected significant evolutionary incompatibilities between conserved single-copy genes, as well as a clear central evolutionary tendency involving 62 gene families that occur as both single and multiple-copies across genomes.

### Methodology

Orthogonal Distance Regression (ODR) is an errors-in-variables regression method that accounts for measurement errors in both explanatory and response variables (Boggs et al. 1987), instead of attributing all errors in the expected values exclusively to the response variable, as performed by Ordinary Least Squares (OLS). While OLS regressions seek to minimize the sum of squared residuals of the response variable, ODR minimizes the sum of squared residuals from each data point obtained by the combination of explanatory and response variables. Novichkov et al. (Novichkov et al. 2004) assessed the compatibility between the evolutionary history of genes with a reference genomic evolutionary history using Pearson’s *r* and estimates of an OLS regression’s intercept. The latter extra step when compared to other implementations using Pearson’s *r* (Ramani and Marcotte 2003; Izarzugaza et al. 2008; Gueudré et al. 2016) is required to infer that datapoints not fitting a regression line through zero are caused by HGT. The approach proposed by Novichkov et al. requires two key assumptions that restrict the general applicability of evolutionary assessments of empirical datasets: 1) there must exist a reference history to which gene histories are compared; and 2) there are no errors in reference distances between genomes.

The approach described here is based on ODR. Its modelling of errors within both assessed variables decreases the necessity of comparing gene family pairwise distances against a well-established reference. Where *a priori* there is no clear separation between explanatory and response variables, errors-in-variables approaches (e.g., ODR) are better suited to compare pairwise evolutionary distances between two gene families. Independently weighing datapoints based on their residuals from an initial regression line provides a framework less susceptible to underestimating overall evolutionary similarities due to few homologs with strong signal incompatibility. Our implementation uses a wODR model with regression line through the origin by setting the Y-axis intercept to zero, which avoids overfitting the linear regression model to the detriment of coherent evolutionary assumptions.

### Algorithm explanation

Our *I_ES_* implementation performs an initial wODR with pairwise comparisons including all gene family representatives within each genome. For each genome, all pairs containing one homolog from each assessed gene family are evaluated, and unique pairs that minimize the sum of squared residuals are reported. As exemplified in Fig. 1, a hypothetical *gene_1_* occurs exclusively as single copy across 10 genomes (Fig. 1, tree_1_), while *gene_2_* has an extra copy within genome *J* (Fig. 1, tree_2_). To identify which copy of *gene_2_* in *J* (i.e., *j_1_* or *j_2_*) better represents their shared evolution we compare *gene_1_* pairwise distances involving *j* with *gene_2_* pairwise distances involving *j_1_* and *j_2_*. Consequently, to do that we must duplicate *J*’s rows and columns in matrix_1_ to match matrix_2_ dimensions (Fig. 1, matrix_1_). The scatter plot in Fig. 1 highlights pairwise distances involving *j_1_* in blue and *j_2_* in red, and as shown by the fitted wODR regression, *gene_2_* pairwise distances involving *j_1_* fits better to the expected linear association between matrix_1_ and matrix_2_ than pairwise distances involving *j_2_*. The smallest sum of residuals obtained by the *j_1_* homolog of *gene_2_* correctly pairs it with *J*’s *gene_1_* homolog, while *j_2_*’s *gene_2_* homolog is likely a product of HGT from a shared common ancestor of *A* and *B*. When both gene families occur in multiples within the same genome, all pairs of unique loci are reported. Once best matching genes from each gene family are paired, or if both occur exclusively as single copy, a final wODR is performed using paired homologs from each gene family and Y-axis intercept equal to zero. wODR is performed through the SciPy (Virtanen et al. 2020) API of ODRPACK (Boggs et al. 1989). Initial weights of pairwise distance are estimated as the inverse of residuals obtained from geometric distance regression with intercept equal to zero and slope equal to *S_Y_/S_X_*, where *S_Y_* and *S_X_* are standard deviations from the regressed distance matrices. Our method’s capability to automatically pair copies of duplicated genes who best reflect the shared history between two gene families vastly expands the scope of datasets fit to general evaluation of evolutionary signal compatibilities. The presence of multiple gene copies within a genome constitutes a key bottleneck to methods commonly used to assess the similarity of evolutionary histories. Tree-based evolutionary distance assessment algorithms are not generally capable of pairing genes between two gene families when at least one family contains multiple gene copies within genomes (Stamatakis 2006; Nguyen et al. 2015; Gori et al. 2016; Huerta-Cepas et al. 2016). While Pearson *r* implementations either rely on multiple tests (Gertz et al. 2003; Ramani and Marcotte 2003; Izarzugaza et al. 2008) or on predicting structural interaction between gene products (Gueudré et al. 2016).

**Fig. 1.**
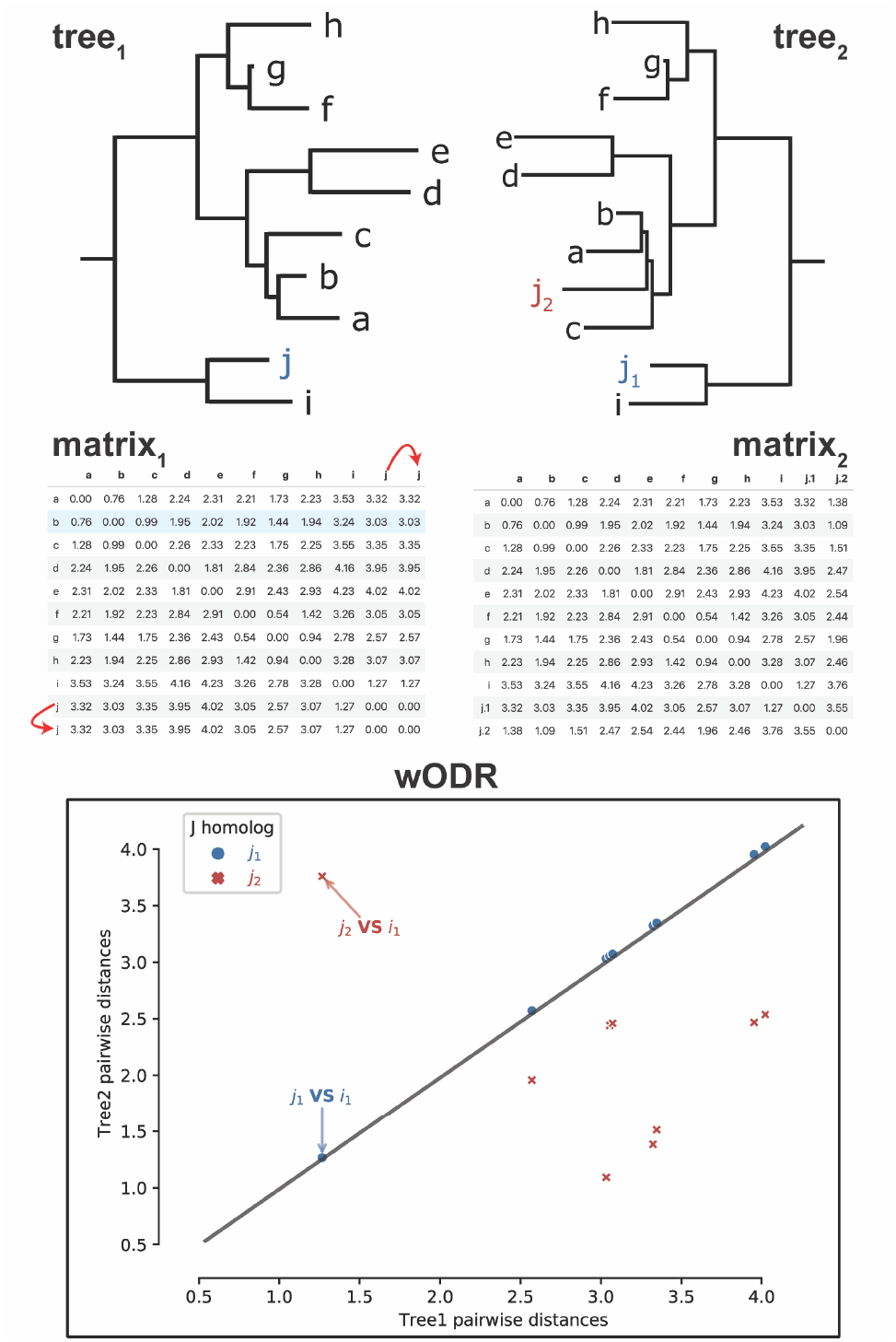
Steps for *I_ES_* estimation between gene families containing multiple gene copies. *tree_1_* and *tree_2_* are phylogenetic trees of two hypothetical gene families, *gene_1_* and *gene_2_*, respectively. *matrix_1_* and *matrix_2_* contain pairwise evolutionary distances between taxa from their respective gene families. The red arrows in *matrix_1_* highlight the duplication of pairwise distances involving the *j* homolog of *gene_1_* necessary to match dimensions of the two matrices. The wODR scatterplot displays the linear relationships between distances from both gene families, and highlights distances related to the *j_1_* homolog of *gene_2_* in blue and related to the *j_2_* homolog in red. Arrows also highlight pairwise distances homologs in genomes *J* and *I* from both gene families.

Given that regression models only account for data points equally represented in both assessed variables, gene losses and duplications are not directly accounted for when comparing evolutionary histories through wODR. To incorporate unequal genomic occurrence between gene families to our proposed measurement of evolutionary similarity, the wODR Coefficient of Determination, i.e. *R*_2_, is adjusted by the Bray-Curtis Index (*I_BC_*).*I_BC_* is defined as 1-*D_BC_*, where is the *D_BC_* is Bray-Curtis Dissimilarity (Bray and Curtis 1957) calculated from absolute genome counts in each gene family. From hereon we will refer to the wODR *R*_2_ × *I_BC_* product as *I_ES_*. Supplementary Fig. S12 demonstrates wODR *R*_2_ overestimation caused by the decrease in taxa overlap between two gene families, as well as the importance of *I_BC_* adjustment. One random taxon was removed from each of two simulated gene families whose evolutions diverge by five Subtree Prune and Regraft (SPR) transformations. As the set of taxa used during the regression becomes unrepresentative of underlying evolutionary processes, estimates based on wODR *R*_2_ tend to artificially increase.

Continuing with the example depicted in Fig. 1, despite *gene1* and *gene2* identical genomic occurrence, their copy numbers diverge within genome *J*, which as mentioned before, arose from a horizontal exchange of *gene2*. To reflect this difference in evolutionary events within gene family histories in the proposed *I_ES_*, the resulting wODR *R*_2_ = 1 is adjusted using an *I_BC_* = 0.95.

### Statistics and data analysis

Pandas Python library (McKinney 2010) was used to manipulate pairwise distance matrices and for generating condensed versions of the matrices submitted to wODR model. Effect size (*f*) hypothesis tests of differences between distributions were obtained using Common Language statistics (McGraw and Wong 1992), and p-value correction for multiple tests was performed using False Discovery Rate implementation in StatsModels Python library (Seabold and Perktold 2010).

Enrichment of gene families within sets of genomes were assessed using hypergeometric tests and p-values corrected with Benjamini-Hochberg’s False Discovery Rate and expressed as q-values.

### Data Simulation

We constructed ten simulated datasets, each one containing 50 trees generated from stepwise random SPR transformations. These datasets were designed to represent sets of trees with similar topologies reflecting a shared evolutionary history, perturbed by both phylogenetic noise and HGT. Each dataset contains one initial random rooted tree with 50 taxa generated by ETE3 (Huerta-Cepas et al. 2016). To obtain the remaining 49 trees, the initial tree (*tree_1*) undergoes a series of 49 consecutive SPR transformations in such a way that *tree_1* differs from *tree_2*, *tree_3*, and *tree_n* by 1, 2, and *n* SPR transformations, respectively. At each SPR the branch leading to the regrafted clade undergoes two transformations to simulate changes in substitution rate after an HGT event. The first transformation multiplies the branch length by a random uniform variable ranging from 0 to 1, simulating at which point during the branch’s history the transfer occurred. The second transformation multiplies by a random gamma distributed variable (α = β = 100), simulating changes in substitution rates in the recipient clade after said transfer. All simulated trees are available in Supplementary Material.

All simulated trees were also used to generate sequence simulations using INDELible (Fletcher and Yang 2009) (Supplementary Material). Phylogenetic trees and pairwise distance matrices were reconstructed using IQTree (Nguyen et al. 2015) using the LG+G model.

### Archaeal empirical dataset

Complete genome sequences of 42 Archaea from the Euryarchaeota phylum and from TACK, DPANN, and Asgardarchaeota groups were downloaded from NCBI GenBank (Supplementary Table S1). Other candidate phyla known from metagenomic as well as remaining members of the DPANN group were not included, as their expected phylogenetic relationships are not as well understood. Clustering of homologous proteins was performed using the orthoMCL (Li et al. 2003) implementation available in the GET_HOMOLOGUES package (Contreras-Moreira and Vinuesa 2013; Vinuesa and Contreras-Moreira 2015). Archaea were selected as the test dataset since the evolutionary relationships between some major groups are well-established, while others remain contested. Furthermore, many sets of archaeal metabolic genes have a strong phyletic dependence (e.g., methanogenesis among Euryarchaeota (Borrel et al. 2013)), therefore facilitating a clear assessment of similarities between evolutionary signals of genes at different phylogenetic distances. Evolutionary similarity comparisons were restricted to homologous groups present in at least 10 genomes.

Pairwise maximum likelihood distances between homologous proteins were generated using IQTree under the LG+G evolutionary model. Phylogenetic trees from clusters of gene families with compatible evolutionary signal (CES) and extended core genome (i.e., single copy and present in at least 35 out of the 42 sampled Archaea genomes) were reconstructed from concatenated multiple sequence alignments using the LG+C60+F+G and individual partitions corresponding to each concatenated gene.

Enrichment of gene functions among CES clusters were performed using StringDB API (Szklarczyk et al. 2019). For each genome, homologs from CES gene families were submitted independently for enrichment assessment. Retrieved protein annotations are available in the Supplementary Material.

### Geodesic and Robinson-Foulds distance calculations

*D_geo_* between single copy gene families (both simulated and real datasets) were calculated using the treeCl Python package (Gori et al. 2016). RF distances between single copy gene families were calculated by ETE3.

## Results and Discussion

### Simulated dataset

Evolutionary histories between simulated gene families were compared to each other using three distinct metrics: RF, *D_geo_*, and *I_ES_*. Results reported by all three approaches successfully identified the monotonic increase in SPR operations from a starting tree (Fig. 2 and Supplementary Fig. S1). Measurements obtained from RF and *D_geo_* approaches, however, frequently overestimated the impact of SPR transformations between two gene families, leading to a fast saturation of dissimilarities between evolutionary histories (Fig. 2B and Fig. 2C). The dissimilarity saturations detected by RF and *D_geo_* measurements occur as they fail to identify the decreasing similarity between two trees separated by more than 15 to 20 SPR transformations, or even 10 SPR in some replicates (simulation replicate #1, Supplementary Fig. S1). Both of these approaches rely on the proportion of compatible bipartitions shared by two trees, which is very susceptible to small changes at deep bipartitions, where changing one single leaf can potentially create fully incompatible bipartition tables.

**Fig. 2.**
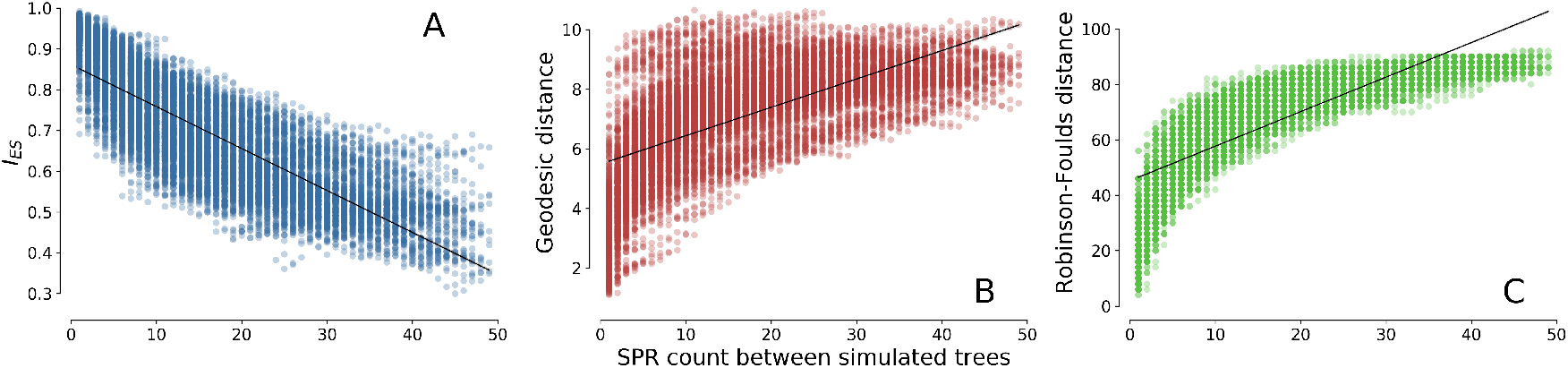
Scatter plot of evolutionary similarity metrics against number of SPR transformations between simulated gene families from all ten replicates. Solid black lines are estimated from OLS regressions between number of SPR transformations and evolutionary similarity metrics. All three scatter plots display the number of SPR transformations between two trees in the X-axis, while varying the evolution similarity metric displayed in the Y-axis. A) displays wODR *R*_2_ between distance matrices of simulated gene families in the Y-axis, B) displays geodesic distances between trees reconstructed from each simulated alignment, and C) displays RF distances estimated from the same trees.

In contrast, *I_ES_* estimates are less susceptible to overestimating the impact of shifts in topology of an underlying tree, displaying a robust linear correlation with the number of SPR transformations between gene families (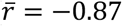, Fig. 2A). The lower level of information assessed by *I_ES_*, pairwise distance matrices instead of dichotomic trees, is less susceptible to dissimilarity saturation, corresponding to a more linear relation between expected and observed changes in evolutionary histories. Furthermore, *I_ES_* is much more efficient, computationally. Tree reconstruction of the alignment simulated from *tree_1* of the first simulation replicate (50 taxa and 500 sites with no indels) under LG+G model by IQTree took 80 seconds in a single thread, while the computation of the pairwise distance matrix for the same alignment took 1.713 seconds. Both computations were performed on a 3 GHz Intel Xeon W processor. The difference in computing time of almost 50x, without bipartition support assessment, shows another, practical advantage for assessing evolutionary similarity through *I_ES_* in large datasets.

### Robustness assessment between approaches

The dichotomic pattern in a cladogram is extremely susceptible to uncertainties in phylogenetic reconstruction, combined with the vast tree space available for 50 taxa, causing noise-induced topological variations to be not directly distinguishable from real deviations in evolutionary history (Szöllosi et al. 2013). The simpler information used to estimate *I_ES_* (i.e., pairwise Maximum Likelihood distances) is less prone to such uncertainty, as it bypasses forming hypotheses about the evolutionary relationships between taxa. This assumption is corroborated by pairwise comparisons within bootstrap replicates, where *I_ES_* correctly detected replicates as such, i.e., virtually identical to each other, while RF and *D_geo_* measures failed to identify the common nature of bootstrap replicates. In addition to its accurate predictions, *I_ES_* consistently displayed very little variance within its estimates between bootstrap replicates.

Each alignment of simulated sequences was used to generate 10 bootstrap replicates. Pairwise comparisons between 10 bootstrap replicates summed up to 45 comparisons within a single alignment. Given that we simulated a total of 500 alignments, we assessed 22,500 pairwise comparisons between bootstrap replicates across all simulated datasets.*I_ES_* values correctly identified bootstrap replicates as sharing virtually identical evolutionary histories, 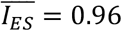, and did so very consistently (CV = 1.39%, where CV stands for Coefficient of Variation). Despite successfully identifying increasing evolutionary changes between simulated trees, RF distances inconsistently predicted similarities between histories of bootstrapped trees (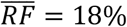 and CV = 57.45%), as variations in bootstrapped alignments caused small perturbations in reconstructed tree topologies, and subsequent underestimation of evolutionary similarity between bootstrap replicates. While *D_geo_* estimates are more likely to overestimate small differences between trees than RF, and consequently are more prone to saturation, geodesic distance estimates displayed substantially less variation than RF (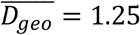 and CV = 14.97%). *D_geo_* cannot be directly normalized or translated to a proportion of incompatibility between trees.

### Evolutionary similarities within archaeal gene families

In order to test *I_ES_* performance when estimating shared phylogenetic signal in an empirical set of gene families, we evaluated 1,322 families of homologous proteins assembled from annotated CDSs extracted from 42 archaeal genomes (Supplementary Table S1). This empirical dataset contains conserved and accessory gene families with different sizes due to gene losses, duplications, and transfers.

*I_ES_* was estimated for all pairwise combinations of gene families present in at least 10 genomes, with 2,142 out of 748,712 comparisons having *I_ES_* values of at least 0.7. Pairs of gene families with an *I_ES_* ≥ 0.7 were added as nodes to a weighted network with its estimated *I_ES_* value as an edge connecting both gene families. In total 419 unique archaeal gene families were added to the network, while the remaining 903 gene families did not display any *I_ES_* ≥ 0.7 with other gene families. A 0.7 threshold was selected based on early applications of Pearson’s *r* to evolutionary distances (Goh et al. 2000; Pazos and Valencia 2001). The resulting evolutionary similarity network (Fig. 3) is heavily imbalanced, with just 11% of nodes involved in 50% of network edges. The majority of gene families (68%) did not display any *I_ES_* above the 0.7 threshold with other gene families, suggesting a general incompatibility between evolutionary signals, or lack thereof to detect its compatibility with others. However, the high edge concentration within just a few nodes suggests a strong central signal (Puigbò et al. 2009) present among few gene families, from which the evolutionary trajectories of others have diverged. Similarities between evolutionary signals, as estimated by *I_ES_*, are strongly associated with genomic linkage (*p* = 6.18 ^-75^ and *f* = 0.86). Gene families frequently occurring in each other’s genomic vicinity (i.e., fewer than 10,000 bp apart in at least 21 genomes) displayed significantly greater *I_ES_* relative to pairs of gene families that were further apart (i.e., more than 100,000 bp apart in at least 21 genomes) (Fig. 4a).

**Fig. 3.**
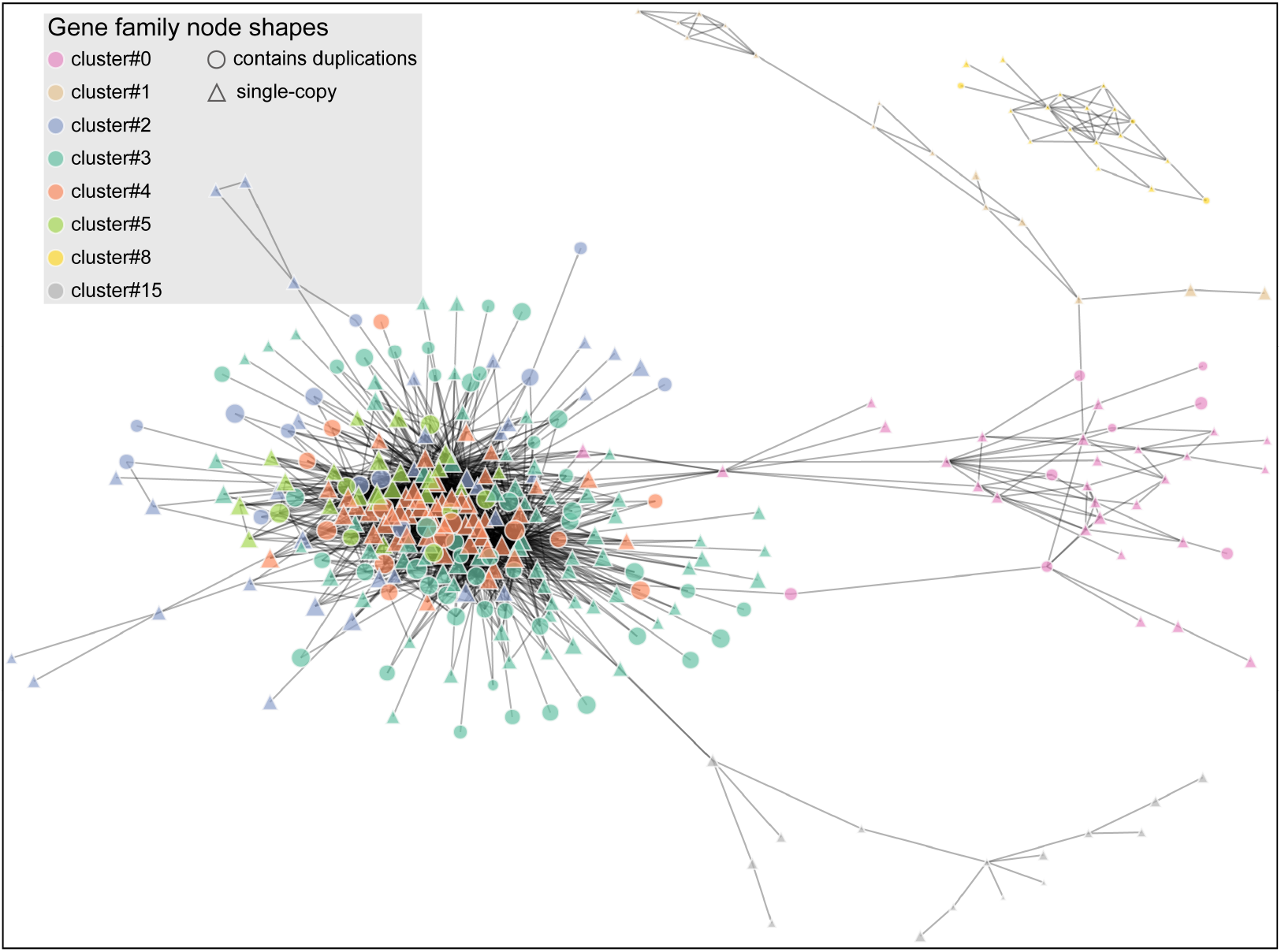
Compatible evolutionary signal network, each node represents a gene family, and edges connecting nodes represent shared evolutionary history (*I_ES_* ≥ 0.7). Nodes in the same colors are identified as undergoing similar evolutionary trends by Louvain community detection. Triangular nodes represent single-copy genes, and circular ones are gene families containing gene duplications. Clusters of CES gene families with less than ten members are not represented.

**Fig. 4.**
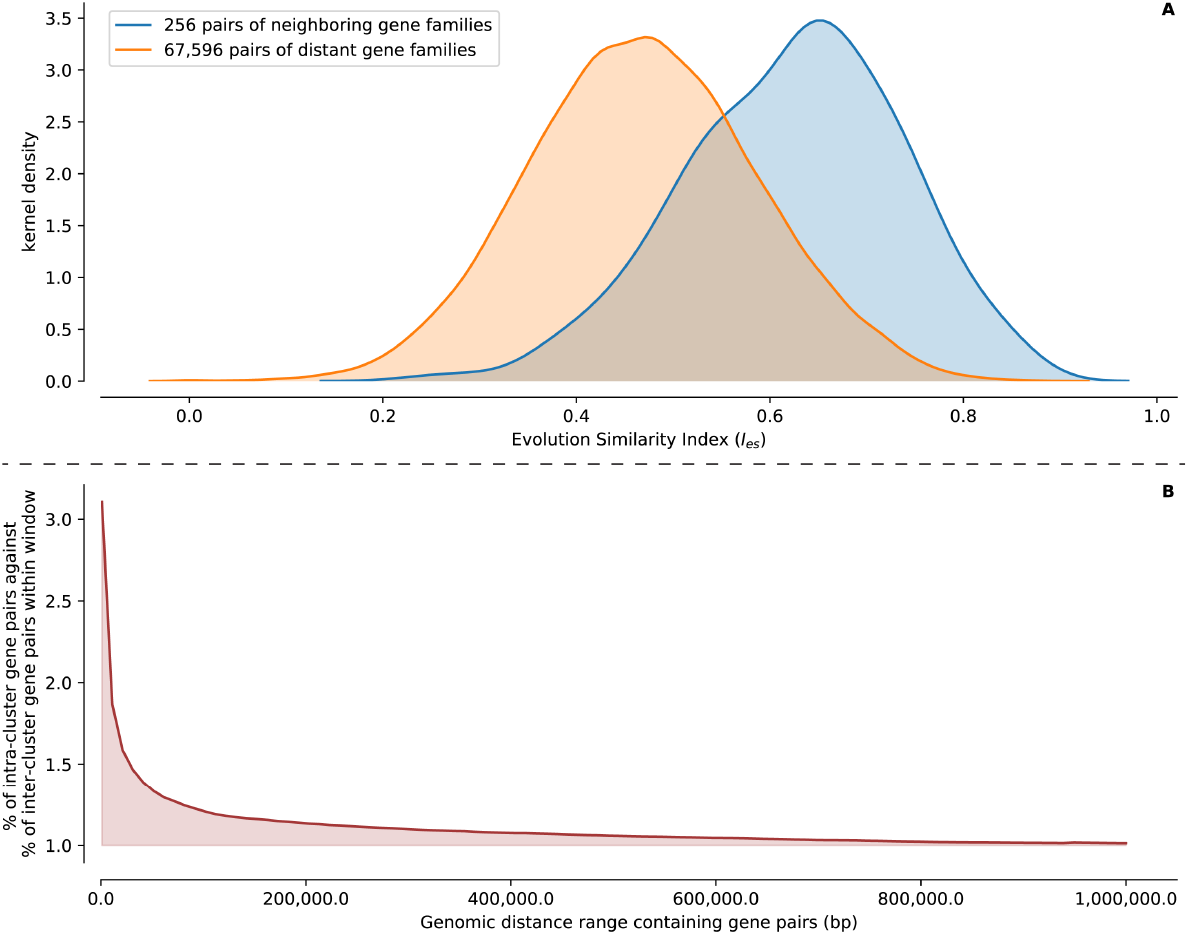
A) distributions of *I_ES_* between pairs of genes within 10,000 bp of each other in blue and between pairs of genes apart by at least 100,000 bp in orange. Neighboring gene pairs displayed significantly more similar evolutionary signals than non-neighboring gene pairs. B) ratio between the proportion of gene pairs intra and inter CES clusters, Y-axis, occurring within genomic windows, X-axis. 100 window sizes were assessed ranging from 1,000 bp to 1,000,000 bp.

### Clusters of gene families with compatible evolutionary signal

Evolutionary trends shared across gene families were assessed using Louvain community detection (Blondel et al. 2008), reporting 41 CES clusters, of which 25 comprise only two gene families and eight major clusters contain ten or more similarly evolving gene families (Fig. 6). Although the assumption of compatibility between evolutionary signals based on *I_ES_* estimates and the clustering process are agnostic of specific shared evolutionary events or their causes, each of these clusters is expected to comprise gene families sharing common evolutionary trends and paths. These shared evolutionary trends are corroborated by the very pronounced association between genomic linkage and estimated compatibility between evolutionary signals. Across small nucleotide distances between loci, linkage is a strong predictor of CES relations between genes, but its predictive power rapidly decreases as the number of nucleotides between two given loci increase (Fig. 4b), displaying a linear log-log relationship (Supplementary Fig. S2). Comparisons between intra- and inter-cluster genomic linkage showed that the proportion of CES genes within 1,000 bp of each other is three times the proportion of non-CES genes within the same window. Increasing the surveilled genomic window decreases the difference between proportions; within a 10,000 bp window, the ratio of CES genes is reduced to 1.8 the ratio of non-CES, and at a 100,000 bp window this difference in proportions falls to 1.2 (Fig. 4b).

Among the eight CES clusters with ten or more gene families, four are comprised of mostly core genes, and four are composed of mostly accessory genes (Fig. 6). The four CES clusters of core genes (cluster#2, cluster#3, cluster#4, and cluster#5) are promising candidates for reconstructing the average phylogenetic signal present within sampled Archaea. These four core CES clusters are composed of 102 extended core genes (single copy and present in at least 35 genomes) and 146 broadly distributed gene families present on average in 33 genomes, both as single and multiple copies. CES clusters of accessory genes (cluster#0, cluster#1, cluster#8, and cluster#15 in Fig. 6) include specific archaeal clades, but do not map to well-established phylogenetic relationships; rather, they show polyphyletic gene distributions, likely caused by HGTs and/or gene losses shared by CES gene families. For example, cluster#0 is well represented amongst Euryarchaeota and hyperthermophilic TACK; cluster#15 comprises gene families with shared evolutionary trends mainly occurring within Crenarachaeota and hyperthermophilic Euryarchaeota; CES accessory gene families in cluster#1 and cluster#8 display congruent signals tying methanogenic Euryarchaeota with Thaumarchaeota and Asgardarchaeota, respectively. Besides the eight CES clusters with ten or more gene families, the CES network community detection yielded 34 other clusters containing between two and nine gene families (Supplementary Material). These 34 CES clusters contain a total of 88 gene families, whose degree centralities are much smaller than the 331 gene families within the eight major CES clusters (averages of 1.36 and 17.79, respectively).

CDSs from 21 out of 42 sampled genomes have functional annotation available in StringDB (Supplementary Material), and through its API we identified annotated KEGG Pathways enriched within homologs of CES gene families from each genome. In the dendrogram and heatmap depicted in Fig. 5 we clearly identify two sets of opposing CES clusters of gene families: accessories (top three rows) and core (bottom four rows), and their associations with genetic information processing and metabolism KEGG Pathways (indicated by column color in the top row). All four CES clusters of core gene families are enriched with KEGG Pathways related to genetic information processing (e.g., Ribosome, DNA replication, and Aminoacyl-tRNA biosynthesis). CES clusters of accessory gene families on the other hand tend to be enriched with KEGG Pathways related to metabolism (e.g., Methane metabolism, Microbial metabolism in diverse environments, and Biosynthesis of antibiotics in Fig. 5). It is also important to emphasize the opposite pattern of enrichment and depletion of KEGG Pathways between clusters of core and accessory genes. KEGG Pathways related to metabolism display minor enrichment signal within CES clusters of core gene families, and KEGG Pathways related to genetic information processing are not enriched within clusters of accessory genes (Fig. 5). CES clusters of accessory genes comprised within cluster#1, whose occurrence is restricted to methanogenic Euryarchaeota and Thaumarchaeota, are enriched for methane metabolism within six genomes. Similarly, gene families from cluster#8, restricted to methanogenic Euryarchaeota and Asgardarchaeota, are also enriched in methane metabolism in five genomes (Fig. 5).

**Fig. 5.**
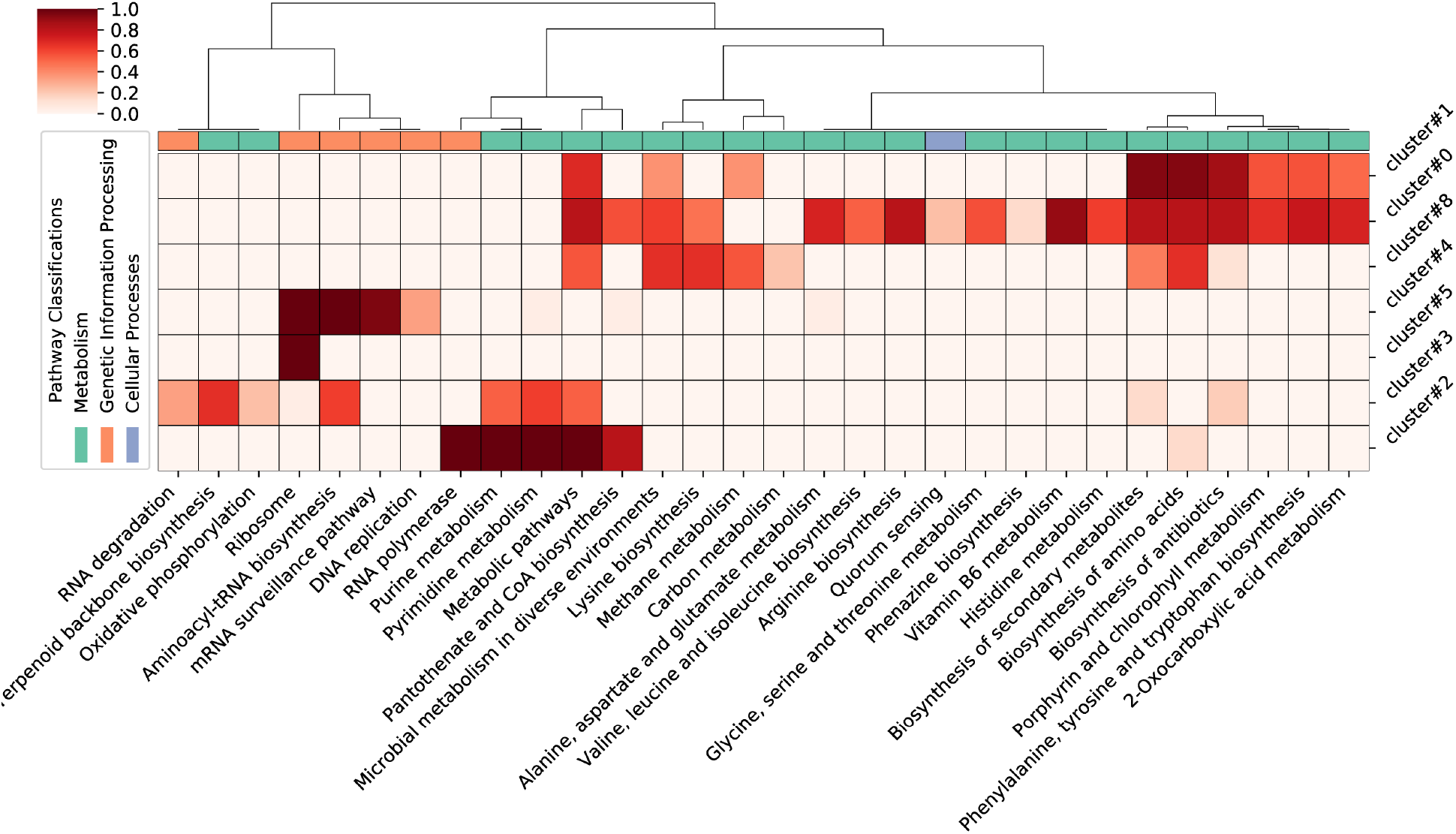
Heatmap of enriched KEGG Pathways, columns, within CES clusters of gene families, rows. Shades of red represent the proportion of genomes with detected KEGG Pathway enrichment within its homologs of CES gene families. Columns and rows were clustered using complete linkage and correlation coefficients. KEGG Pathways enriched in less than 10% of genomes in which CES genes occur are not reported. Cluster#15 did not report significant enrichment of KEGG Pathways.

### Compatible evolutionary signal clusters and possible vertical evolutionary signals

Phylogenies generated from extended core genomes are generally used as reasonable proxies of the species-tree phylogeny, given the assumption that these genes are less likely to undergo HGT between distantly related groups. However, an extended core phylogeny may not represent the species tree for several reasons, including systematic biases in phylogenetic reconstruction due to shared compositional bias, or strong biases in HGT partners among sets of genes. Nevertheless, the extended core tree can still be used as an adequate representation of the consensus evolutionary signal detected in the sampled archaeal genomes, the closest thing we have to the simple “null hypothesis” of a shared history due to vertical inheritance. The 102 genes composing the extended core genome are not equally distributed across CES clusters (Fig. 3), e.g., cluster#4 contains the greatest number of extended core genes, 44 out of 62 gene families, followed by cluster#3 with 27 among its 111 gene families. The split of the extended core genome into four distinct major CES clusters (Fig. 3) suggests differing sets of HGTs among core genes, creating conflicting evolutionary histories between genes from different clusters (Fig. 7). Closeness centrality measures 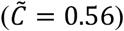 and node strength corrected by cluster size 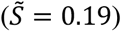 suggest that cluster#4 gene families share stronger and more cohesive evolutionary trends than gene families from other clusters (Supplemental Fig S3). Therefore, cluster#4 contains the set of genes that may be best able to approximate the average evolutionary signal detected among sampled Archaea as well the most compatible to each other. Cluster #4’s evolutionary history is also the most similar to that inferred from the extended core genome (Fig. 6 and Fig. 7).

**Fig. 6.**
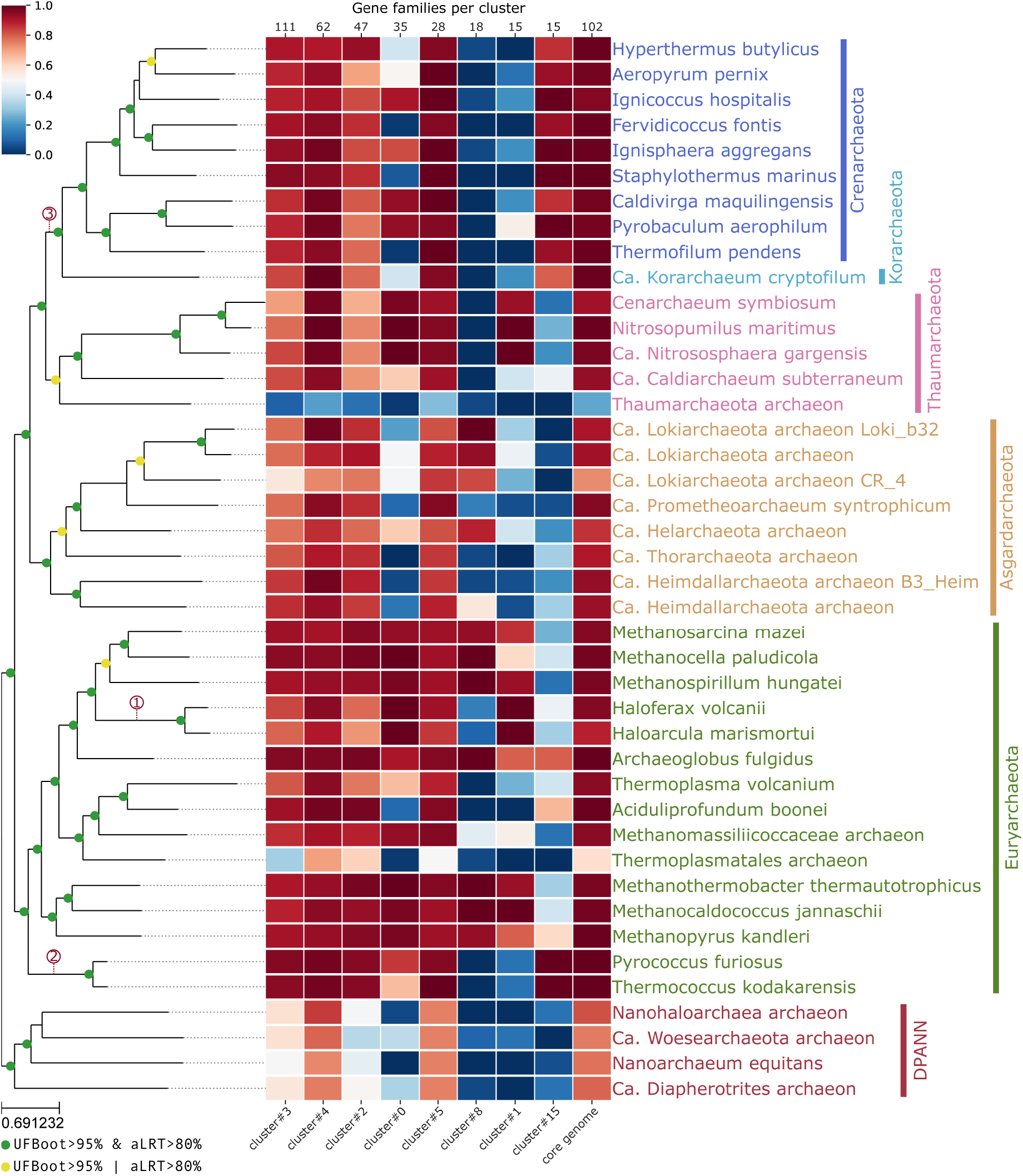
Phylogenetic tree of Archaea reconstructed from 62 genes within CES cluster#4. The phylogeny was obtained using LG+F+G+C60 evolutionary model from IQTree and each gene had its parameters independently estimated according to parameter “-sp”. Bipartition supports were estimated using UFBoot and aLRT, each with 1,000 replicates, and bipartitions well supported by both methods are colored in green (UFBoot ≥ 95% and aLRT ≥ 80%), while bipartitions well supported by a single method are colored in yellow. Red dotted lines indicate Nanohaloarchaea (1) and Nanoarchaea (2 and 3) placements reconstructed within phylogenies containing a single DPANN genome at a time. Despite the lack of outgroups to Archaea within our sample the tree is rooted in DPANN for the sake of visualization. The associated heatmap reflects the representation of gene families within CES clusters amongst archaeal genomes.

**Fig. 7.**
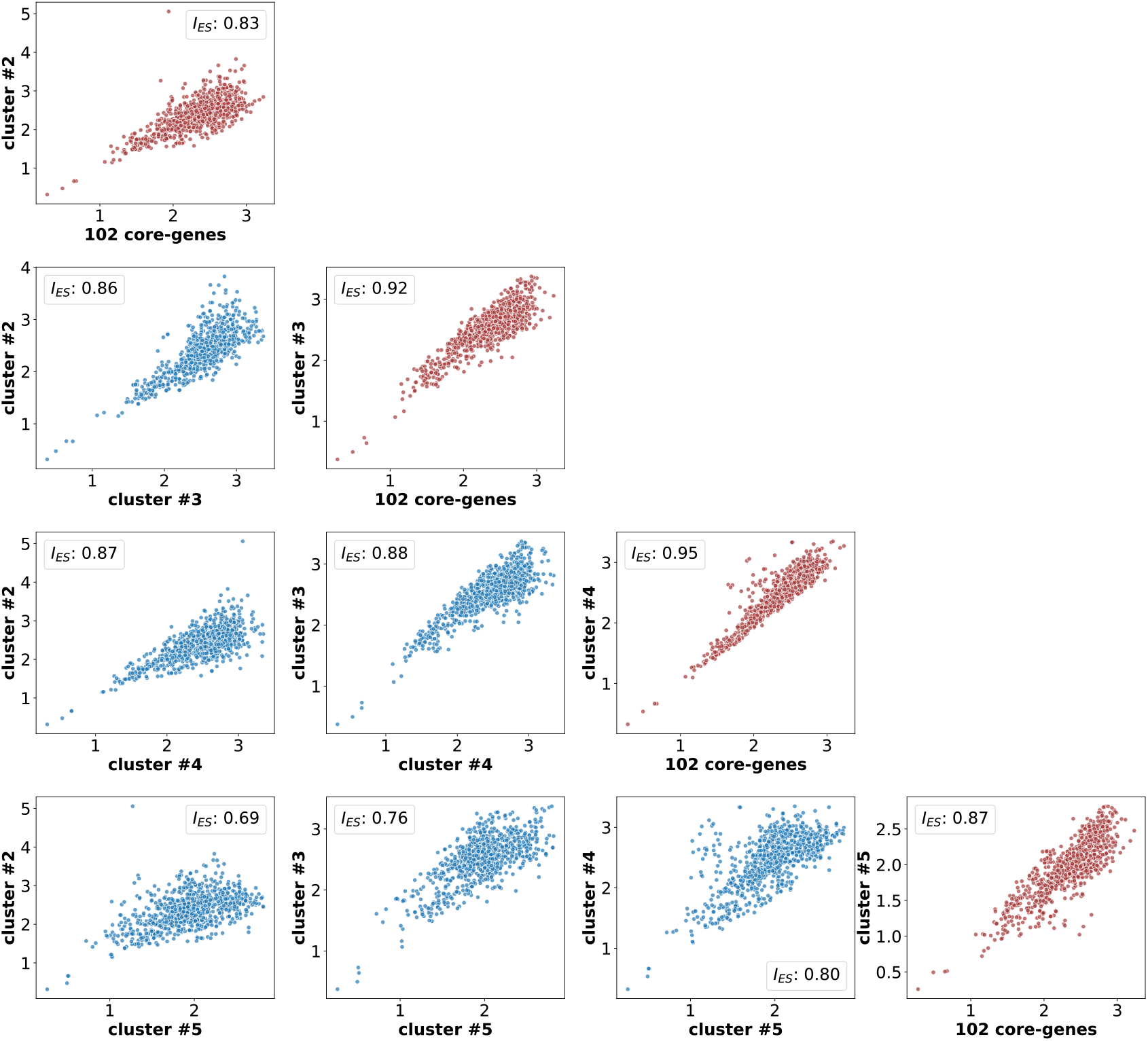
Scatter plots of pairwise evolutionary distances reconstructed from each widely distributed gene family versus each other in blue. And in red, scatter plots of pairwise evolutionary distances reconstructed from each widely distributed gene family versus pairwise evolutionary distances reconstructed from 102 extended core gene families. Similarities between evolutionary histories of pairs of CES clusters, and between CES clusters and extended core genes were estimated by *I_ES_*.

While binning genes with distinct evolutionary histories permits phylogenetic reconstructions less likely to be subject to spurious signals arising from the averaging of conflicting evolutionary signals, the resulting phylogenies remain susceptible to phylogenetic reconstruction artefacts. For example, since *I_ES_* is not estimated from phylogenies but from pairwise evolutionary distances, we do not expect it to be subject to Long Branch Attraction (LBA) artefacts. Nevertheless, phylogenies reconstructed from sets of genes with high *I_ES_* between each other are as susceptible to LBA as any other dataset. By clustering gene families with high *I_ES_* we are able to minimize conflicting evolutionary signals between concatenated genes, improving the robustness of the phylogenetic signal. Despite its robustness to phylogenetic artifacts, *I_ES_* estimates are still affected by sampling biases. The overrepresentation of specific taxonomic groups can lead to underestimating deviations in the evolutionary history of less represented groups.

LBA is a frequently invoked in discussions of archaeal phylogeny, specifically with regard to the phyletic status of the DPANN group (Brochier-Armanet et al. 2011; Raymann et al. 2014; Petitjean et al. 2015; Williams et al. 2015; Feng et al. 2019). Regardless of the set of genes used for phylogenetic reconstruction, extended core genome or any of the four CES clusters of core genes, all resulting trees depicted a well-supported DPANN clade composed of Nanohaloarchaea archaeon, *Ca*. Woesearchaeota archaeon, *Nanoarchaeum equitans*, and *Ca*. Diapherotrites archaeon. To assess the impact of LBA in reconstructing DPANN, we generated phylogenies from the CES clusters of core genes while using a single DPANN taxon at a time. Each DPANN taxon showed distinct evolutionary trends across all CES clusters of genes, suggesting both a more complex extended core gene history for these genomes, and that the initial monophyletic grouping of DPANN in each case was, in fact, artifactual (Supplementary Material).

Cluster#4 phylogenies individually testing the position of each DPANN taxa placed Nanohaloarchaea sister to Halobacteria, with Nanohaloarchaea+Halobacteria being sister to Methanomicrobia. Cluster#3 also reported Nanohaloarchaea sister to Halobacteria, but with both nested within Methanomicrobia, rendering this group paraphyletic. This placement for Nanohaloarchaea has been previously proposed by Narasingarao et al. 2012; Zhaxybayeva et al. 2013; Petitjean et al. 2015; and Feng et al. 2019. The uncertain placement of Nanoarchaea has also been topic of investigation (Huber et al. 2002; Brochier et al. 2005). Interestingly, each CES cluster recovered a different placement for Nanoarchaea: sister to Euryarchaeota (cluster#2), sister to Korarchaeota+Crenarchaeota (cluster#3), sister to Korarchaeota (cluster#4), and sister to Thermococcales (cluster#5). One of the most accepted Nanoarchaea placements is as sister to Thermococccales (Brochier et al. 2005; Urbonavičius et al. 2007; Dutilh et al. 2014), which in our analyses was recovered only by cluster#5 (Supplementary Material).

While our tests further support that the monophyly of DPANN is likely due to LBA, we did not detect a significant LBA effect for Woesearchaeota and Diapherotrites. Except for Diapherotrites placing between Class I and II methanogens in cluster#2, phylogenies from all four clusters proposed both taxa grouping together as sister to Euryarchaeota, assuming an archaeal root between TACK+Asgard and Euryarchaeota (Supplementary Material). The disparate placements of DPANN members within trees from CES clusters also suggests that, in addition to LBA, the DPANN clade from the extended core genome phylogeny is further impacted by the heterogeneity of the phylogenetic signal. This may not only produce a “signal averaging” effect favoring a monophyletic DPANN deeper in the archaeal tree, but may also be a contributing factor to the LBA artifact itself. Heterogeneity among combined phylogenetic signals is likely to increase the estimated branch length, as the incorrect assumption of a single underlying phylogeny will lead to more homoplastic sites.

Assuming that a given set of genomes constitutes a monophyletic clade, it is also reasonable to expect a certain number of gene families to be overly represented within the clade and not readily available to genomes outside the clade. Regardless of the driving force behind the enrichment of gene families within a clade, inheritance from a common ancestor or biased HGT (Andam et al. 2010; Andam and Gogarten 2011), we identified 80 gene families enriched within TACK genomes and 111 within Euryarchaeota (*q* ≤ 0.05). In contrast to the well accepted TACK and Euryarchaeota clades, we did not detect any number of genes enriched within the four sampled DPANN genomes, providing phylogenetic independent evidence against its monophyly. Complementarily, and in support of a Nanohaloarchaea+Halobacteria clade, we identified 78 genes present in Nanohaloarchaea archaeon enriched within the three Nanohaloarchaea and Halobacteria genomes. When compared to well accepted clades containing a similar number of sampled genomes to DPANN, 456 gene families are enriched within the five genomes within Thaumarchaeota. While the differing degrees of physiological, metabolic, and genetic diversity within these groups certainly influence the number of shared gene families, it remains striking that this particular signal of shared ancestry is conspicuously lacking in DPANN.

### Common and distinct evolutionary trends between CES clusters

Among CES clusters of core gene families, cluster#4 and cluster#5 are most evenly represented across archaeal groups, while cluster#2 and cluster#3 are poorly distributed among DPANN (Fig. 6). All four CES clusters of core gene families have low frequency within Thaumarchaeota archaeon SCGC AB-539-E09, and only gene families from cluster#2 and cluster#4 are present in any substance in Thermoplasmatales archaeon SCGC AB-539-N05. All four clusters display very similar overall phylogenies calculated from concatenations of genes within each cluster, varying mainly within the organization of Euryarchaeota (Fig. 6 and Supplementary Material). All four CES clusters of core genes reconstructed the monophyly of Euryarchaeota, with the exception of cluster#2, which placed *Pyrococcus furiosus, Thermococcus kodakarensis, Methanocaldococcus jannaschii, Methanothermobacter thermautotrophicus*, and *Methanopyruus kandleri* together as sister to Asgardarchaeota+TACK. Only cluster#4 recovered the monophyly of Methanomicrobia as sister to Halobacteria, with the other three CES clusters placing Halobacteria within Methanomicrobia.

All four core CES clusters robustly identified Asgardarchaeota as sister to TACK (Fig. 6), with small variation in the Asgardarchaeota phylogeny, and cluster#5 placed Korarchaeota at the base of the TACK super-phylum. When assessing all-versus-all *I_ES_* between CES clusters of core genes, the evolutionary signal detected from cluster#4 is the least dissimilar to the other three (Fig. 7). This shortest path from cluster#4’s evolutionary trajectory to others suggests that cluster#4 best approximates the average archaeal evolutionary history (Fig. 7). In general, the overall high *I_ES_* estimates between core CES clusters suggest that despite composing distinct clusters, gene histories between clusters are generally congruent, with deviations reflecting small divergences potentially representing genes with specific sets of reticulate histories.

Phylogenetic trees obtained from accessory gene families in cluster#0, cluster#1, cluster#8, and cluster#15 reconstructed all represented archaeal phyla as monophyletic (except for *P. furiosus* in Euryarchaeota in cluster#0, Supplementary Fig. S5), suggesting a shared common origin of accessory genes from each CES cluster by all genomes from the same phylum. Although the monophyly of archaeal phyla within trees of CES clusters of accessory genes does not permit an accurate prediction of the directionality of possible inter-phyla HGTs, intra-phylum distances congruent to the supposed vertical inheritance signal can be used to evaluate inter-phylum distances under a wODR model (Supplemental Fig. S6, S8, S10, and S11). When compared to pairwise distances expected from vertical inheritance, inter-phylum distances that are significantly shorter than estimates obtained from intra-phylum distances may be attributed to HGT acquisition by one of the phyla in question. For each CES cluster of accessory genes, we assessed wODR of its pairwise distances against the vertical evolution estimated from cluster#4.

When comparing pairwise distances obtained from cluster#1 against cluster#4, distances between Euryarchaeota and Thaumarchaeota are consistently placed below the estimated regression line (Supplementary Fig. S6 and S7). This suggests that cluster#1 genes were horizontally transferred between ancestors of both phyla, causing shorter evolutionary distances between phyla than expected if their homologs diverged exclusively by vertical inheritance.

Inter-phyla distances between Euryarchaeota and Crenarchaeota obtained from cluster#0 fit the evolutionary rate expected using intra-phylum distances for this CES cluster (Supplementary Fig. S8), suggesting that homologs from both phyla were vertically inherited from a common ancestor. On the other hand, cluster#0 inter-phyla distances involving Thaumarchaeota (Crenarchaeota to Thaumarchaeota and Euryarchaeota to Thaumarchaeota) are shorter than expected from the wODR using intra-phylum distances (Supplementary Fig. S8) and display significantly greater residuals than distances between Crenarchaeota and Euryarchaeota (Supplementary Fig. S9). The absence of cluster#0 genes among Asgardarchaeota and Korarchaeota and the short inter-phyla distances to Thaumarchaeota homologs suggest an extensive loss among missing clades and horizontal acquisition by the thaumarchaeal ancestor from either crenarchaeal or euryarchaeal donors.

Despite the occurrence of accessory genes from cluster#1 and cluster#8 in methanogenic Euryarchaeota (Fig. 6) and the enrichment of methane metabolism pathways (Fig. 5), evolutionary histories of both CES clusters are not related (Fig. 3). Gene families in CES cluster#8 did not display *I_ES_* ≥ 0.7 outside its own cluster, constituting a separate connected component in the CES network depicted in Fig. 3. That said, cluster#8 gene families display shorter Euryarchaeota-Asgardarchaeota distances when compared to cluster#4 distances, but unlike cluster#0 and cluster#1, intra-Asgardarchaeota and intra-Euryarchaeota pairwise distances are not mutually compatible under a single linear regression (Supplemental Fig. S10). The lack of a strong wODR anchor in the form of intra-phyla distances suggests a more complex horizontal exchange history of cluster#8 genes, possibly involving intra-phylum HGTs, which we cannot accurately assess with the dataset used in this study. CES cluster#15 of accessory genes is well distributed among Crenarchaeota, and its intra-phylum pairwise distances are congruent to cluster#4 distances, but their patchy occurrence among Euryarchaeota and Korarchaeota (Fig. 6) does not permit a confident assessment of this cluster’s evolutionary history (Supplementary Fig. S5).

## Conclusions

We have presented *I_ES_*, a new, robust, and efficient method to determine gene families with compatible evolutionary histories that are good candidates to be used in phylogenetic tree reconstructions of a set of organisms. The distance regression basis of our proposed method does not require hypotheses regarding evolutionary relationships between extant and ancestral taxa represented by the branching pattern of phylogenetic trees. Besides significant gains in accuracy of its estimates and computing efficiency, *I_ES_* introduces a new and robust approach to pair members of gene families that best represent their shared evolutionary trends. The strong association between genomic linkage and *I_ES_* estimates within archaeal genomes constitutes independent evidence of the ability of *I_ES_* to recover shared evolutionary histories within empirical datasets.

One major consequence of shared evolutionary trends by gene families is that the set of genomes in which a given gene family occurs should be similar to genomes in which genes of other families with compatible signals occur. Despite similar performances of Pearson’s *r* and wODR *R*_2_ in detecting these trends, *I_ES_* achieves the same result in a more efficient way. The utilization of wODR also imparts more robust statistical support not directly available to previous Pearson’s *r* implementations, whereas the assessment of pairwise distances between taxa provides robustness in the presence of artefacts associated to phylogenetic inference (e.g., Long Branch Attraction). The ability to assess residuals of each datapoint independently also allows for evaluations of specific homologs, a useful tool for HGT detection. *I_ES_* can thus be incorporated into phylogenomics pipelines and used to guide the selection of gene families for more accurate and robust species-tree inference, as well as the detection of meaningful clusters of gene families evolved in shared, yet reticulate, patterns.

By assessing similarities of evolutionary signal between archaeal gene families using *I_ES_* we were able to detect several clusters of shared, distinct gene histories. Phylogenetic reconstruction using concatenated sequences from each of the four major CES clusters of core genes confirmed these distinct evolutionary histories. The phylogeny resulting from CES cluster#4, in particular, recovers a species tree hypothesis consistent with that proposed in several other studies, while using a more empirically-supported selection of gene families that does not presuppose vertical inheritance, an improved alternative to analyses limited to conserved single-copy core genes.

Besides its impact in improving the reconstruction of the Archaeal Tree of Life, CES clusters obtained from *I_ES_* provided key evidence about horizontal exchange between phyla of functionally related genes (Supplementary Fig. S5). For example, given the almost exclusive occurrence of genes from CES cluster#1 among methanogenic Euryarchaeota and Thaumarchaeota, tree-based approaches are not capable to report the possible HGT between both phyla. Separately, intra- and inter-phyla distances obtained from CES cluster#1 are strongly correlated to distances described in CES cluster#4, however the significant placement of inter-phyla distances bellow the wODR line strongly suggests an HGT between ancestors of both phyla.

The method we used to analyze the archaeal gene sets is general and can thus be applied to any other set of genomes. Furthermore, the *I_ES_* implementation described provides a straightforward framework for future improvements and a possible alternative to phylogenetic reconciliation approaches to identify HGT events, as described in Supplementary Figs. S5-S11.

## Acknowledgements

This work was supported by Simons Foundation Collaboration on the Origins of Life Award #339603 and NSF Integrated Earth Systems Program Award #1615426 to GPF. SMS was supported through Geisel School of Medicine at Dartmouth’s Center for Quantitative Biology through a grant from the National Institute of General Medical Sciences of the National Institutes of Health under Award Number P20GM130454. JCS was funded in part by a CNPq senior researcher fellowship.

