## Supporting Figures for "Non-phylogenetic identification of co-evolving genes for reconstructing the archaeal Tree of Life"

Table S1
List of 42 Archaeal genomes download from NCBI GenBank. Columns were extracted from
https://ftp.ncbi.nlm.nih.gov/genomes/genbank/assembly_summary_genbank.txt

| Assembly accession | bioproject | biosample | taxid | Organism name | Infraspecific name | isolate |
|---|---|---|---|---|---|---|
| GCA_000258425.1 | PRJNA91117 | SAMN02603095 | 1163730 | Fervidicoccus fontis Kam940 | strain=Kam940 | |
| GCA_001871415.1 | PRJNA297582 | SAMN04328224 | 1805424 | Candidatus Woesearchaeota archaeon CG1_02_57_44 | | CG1_02_57_44 |
| GCA_002779065.1 | PRJNA362739 | SAMN06659264 | 1974404 | Candidatus Diapherotrites archaeon CG08_land_8_20_14_0_20_34_12 | | CG08_land_8_20_14_0_20_34_12 |
| GCA_000349645.1 | PRJNA168253 | SAMN02261087 | 1198116 | Thermoplasmatales archaeon SCGC AB-539-N05 | | SCGC AB-539-N05 |
| GCA_000015945.1 | PRJNA17449 | SAMN02598390 | 399550 | Staphylothermus marinus F1 | strain=F1 | |
| GCA_003144275.1 | PRJNA383916 | SAMN07236661 | 2012493 | Candidatus Heimdallarchaeota archaeon B3_Heim | | B3_Heim |
| GCA_000145985.1 | PRJNA33361 | SAMN00016987 | 583356 | Ignisphaera aggregans DSM 17230 | strain=DSM 17230 | |
| GCA_000011185.1 | PRJNA206 | SAMD00061089 | 273116 | Thermoplasma volcanium GSS1 | strain=GSS1 | |
| GCA_000025685.1 | PRJNA12524 | SAMN02604027 | 309800 | Haloferax volcanii DS2 | strain=DS2 | |
| GCA_000018305.1 | PRJNA17421 | SAMN00623034 | 397948 | Caldivirga maquilingensis IC-167 | strain=IC-167 | |
| GCA_000009965.1 | PRJNA13213 | SAMD00061071 | 69014 | Thermococcus kodakarensis KOD1 | strain=KOD1 | |
| GCA_003345545.1 | PRJNA406094 | SAMN08287972 | 2053491 | Candidatus Thorarchaeota archaeon | | OWC2 |
| GCA_000200715.1 | PRJNA202 | SAMN02744041 | 414004 | Cenarchaeum symbiosum A | | |
| GCA_000008645.1 | PRJNA289 | SAMN02603244 | 187420 | Methanothermobacter thermautotrophicus str. Delta H | strain=Delta H | |
| GCA_000007065.1 | PRJNA300 | SAMN02603290 | 192952 | Methanosarcina mazei Go1 | strain=Go1 | |
| GCA_000008665.1 | PRJNA104 | SAMN02603985 | 224325 | Archaeoglobus fulgidus DSM 4304 | strain=DSM 4304 | |
| GCA_000007225.1 | PRJNA172 | SAMN02604075 | 178306 | Pyrobaculum aerophilum str. IM2 | strain=IM2 | |
| GCA_000019605.1 | PRJNA16525 | SAMN02598368 | 374847 | Candidatus Korarchaeum cryptofilum OPF8 | | |

| | | | | | | |
|---|---|---|---|---|---|---|
| GCA_005223125.1 | PRJNA383916 | SAMN07236659 | 2012491 | Candidatus Lokiarchaeota archaeon Loki_b32 | | Loki_b32 |
| GCA_000015145.1 | PRJNA208 | SAMN02604082 | 415426 | Hyperthermus butylicus DSM 5456 | strain=DSM 5456 | |
| GCA_000303155.1 | PRJNA60505 | SAMN02603264 | 1237085 | Candidatus Nitrososphaera gargensis Ga9.2 | | enrichment culture Ga9.2 |
| GCA_000017945.1 | PRJNA13914 | SAMN02598324 | 453591 | Ignicoccus hospitalis KIN4/I | strain=KIN4/I | |
| GCA_011364945.1 | PRJNA495098 | SAMN10218972 | 2026747 | Candidatus Heimdallarchaeota archaeon | | SZ_4_bin5.60 |
| GCA_000007185.1 | PRJNA294 | SAMN02603235 | 190192 | Methanopyrus kandleri AV19 | strain=AV19 | |
| GCA_013343275.1 | PRJNA588232 | SAMN13231772 | 2594798 | Nanohaloarchaea archaeon | strain=M3_22 | |
| GCA_000011125.1 | PRJNA211 | SAMD00061092 | 272557 | Aeropyrum pernix K1 | strain=K1 | |
| GCA_000025665.1 | PRJNA38403 | SAMN02598523 | 439481 | Aciduliprofundum boonei T469 | strain=T469 | |
| GCA_000011005.1 | PRJDA20361 | SAMD00060931 | 304371 | Methanocella paludicola SANAE | strain=SANAE | |
| GCA_000270325.1 | PRJDA49157 | SAMD00016619 | 311458 | Candidatus Caldiarchaeum subterraneum | | |
| GCA_000018465.1 | PRJNA19265 | SAMN00000032 | 436308 | Nitrosopumilus maritimus SCM1 | strain=SCM1 | |
| GCA_013375405.1 | PRJNA599172 | SAMN14414634 | 2719382 | Candidatus Helarchaeota archaeon | | CR_Bin_291 |
| GCA_000007305.1 | PRJNA287 | SAMN02604284 | 186497 | Pyrococcus furiosus DSM 3638 | strain=DSM 3638 | |
| GCA_009911715.1 | PRJNA258248 | SAMN02991093 | 1535962 | Methanomassiliicoccaceae archaeon DOK | strain=DOK | |
| GCA_000015225.1 | PRJNA16331 | SAMN02598366 | 368408 | Thermofilum pendens Hrk 5 | strain=Hrk 5 | |
| GCA_008000775.1 | PRJNA557562 | SAMN12405820 | 2594042 | Candidatus Prometheoarchaeum syntrophicum | strain=MK-D1 | |
| GCA_000011085.1 | PRJNA105 | SAMN02603385 | 272569 | Haloarcula marismortui ATCC 43049 | strain=ATCC 43049 | |
| GCA_008080735.1 | PRJNA521734 | SAMN10909897 | 2053489 | Candidatus Lokiarchaeota archaeon | | BC3 |
| GCA_000008085.1 | PRJNA9599 | SAMN02603208 | 228908 | Nanoarchaeum equitans Kin4-M | | |
| GCA_000013445.1 | PRJNA13015 | SAMN02598287 | 323259 | Methanospirillum hungatei JF-1 | strain=JF-1 | |
| GCA_001940655.1 | PRJNA288027 | SAMN04958229 | 1849166 | Candidatus Lokiarchaeota archaeon CR_4 | | CR_4 |

| | | | | | | |
|---|---|---|---|---|---|---|
| GCA_000091665.1 | PRJNA102 | SAMN02603984 | 243232 | Methanocaldococcus jannaschii DSM 2661 | strain=DSM 2661 | |
| GCA_000349625.1 | PRJNA168252 | SAMN02261086 | 1198115 | Thaumarchaeota archaeon SCGC AB-539-E09 | | SCGC AB-539-E09 |

Fig S1

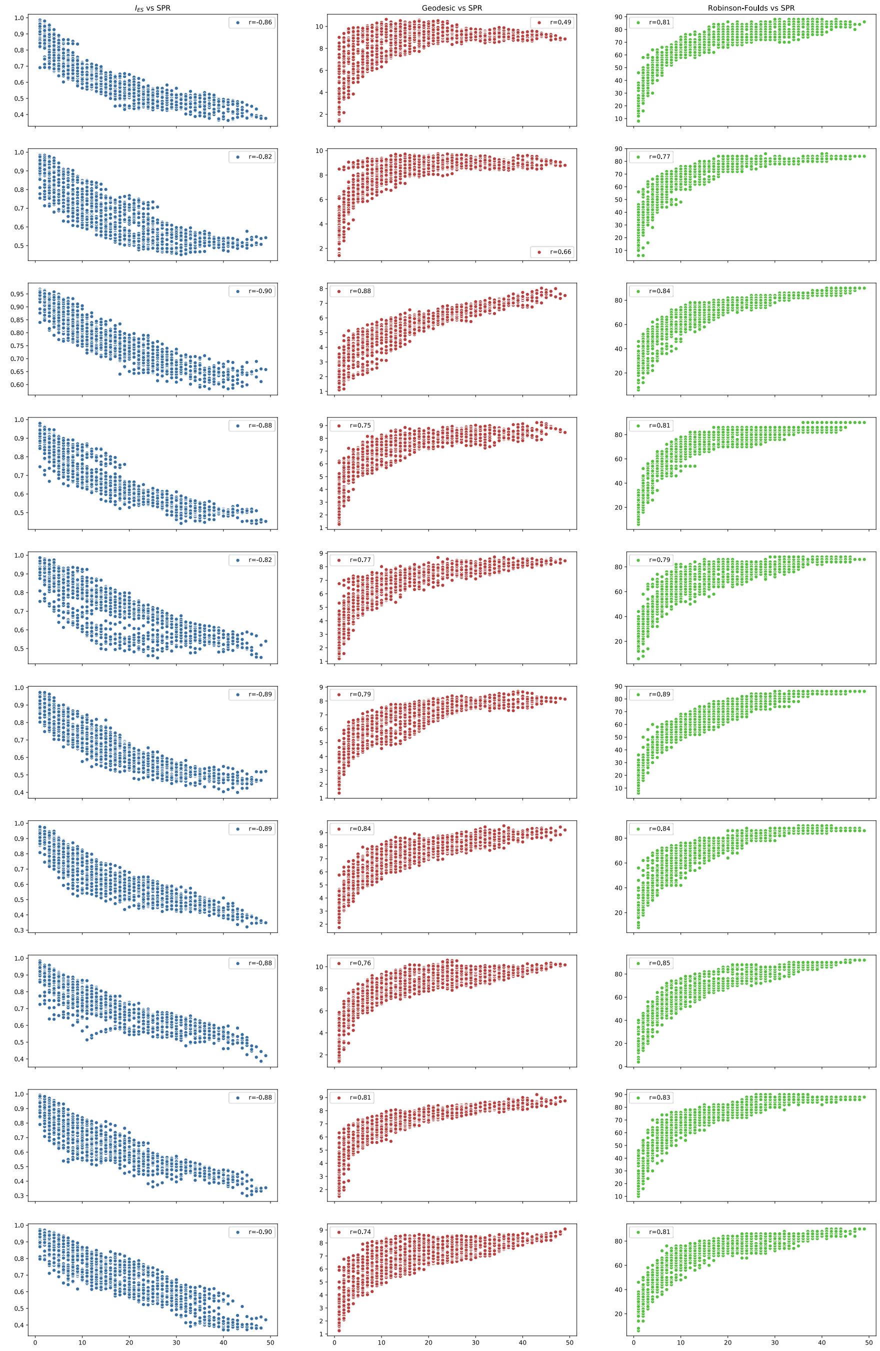

Scatter plot for each simulated tree comparing results from $I_{ES}$ in blue, geodesic distance between trees in red, and Robinson-Foulds in green. All scatter plots display the number of SPR transformations between two trees in the X-axis, while varying the evolution similarity metric displayed in the Y-axis. Pearson correlation coefficients between evolutionary similarity metrics estimated for each simulated dataset and the number of SPR transformations between trees are presented within each scatter plot. The combined data are depicted in Fig. 2.

Fig S2

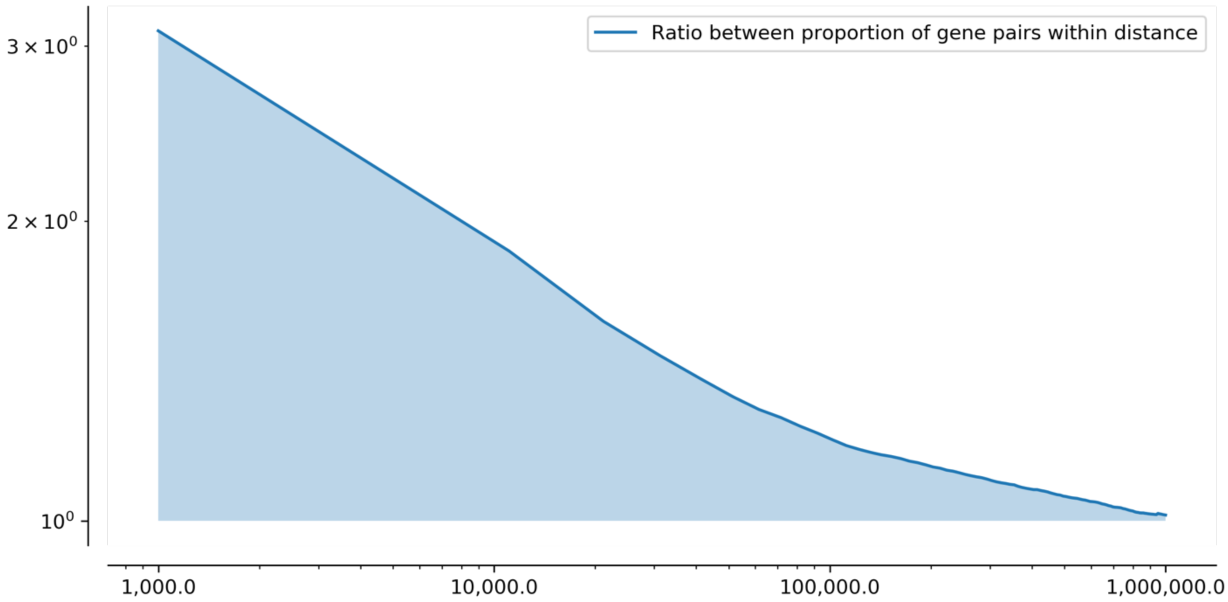

Data from Fig. 4b represented in a log-log scale. Decrease in the proportion of co-evolving over non-co-evolving gene pairs as we increase the surveilled genomic window. The approximation of a linear relationship between log-log scales suggests a power-law decrease in the predictive power of gene co-evolution from genomic linkage.

Fig S3

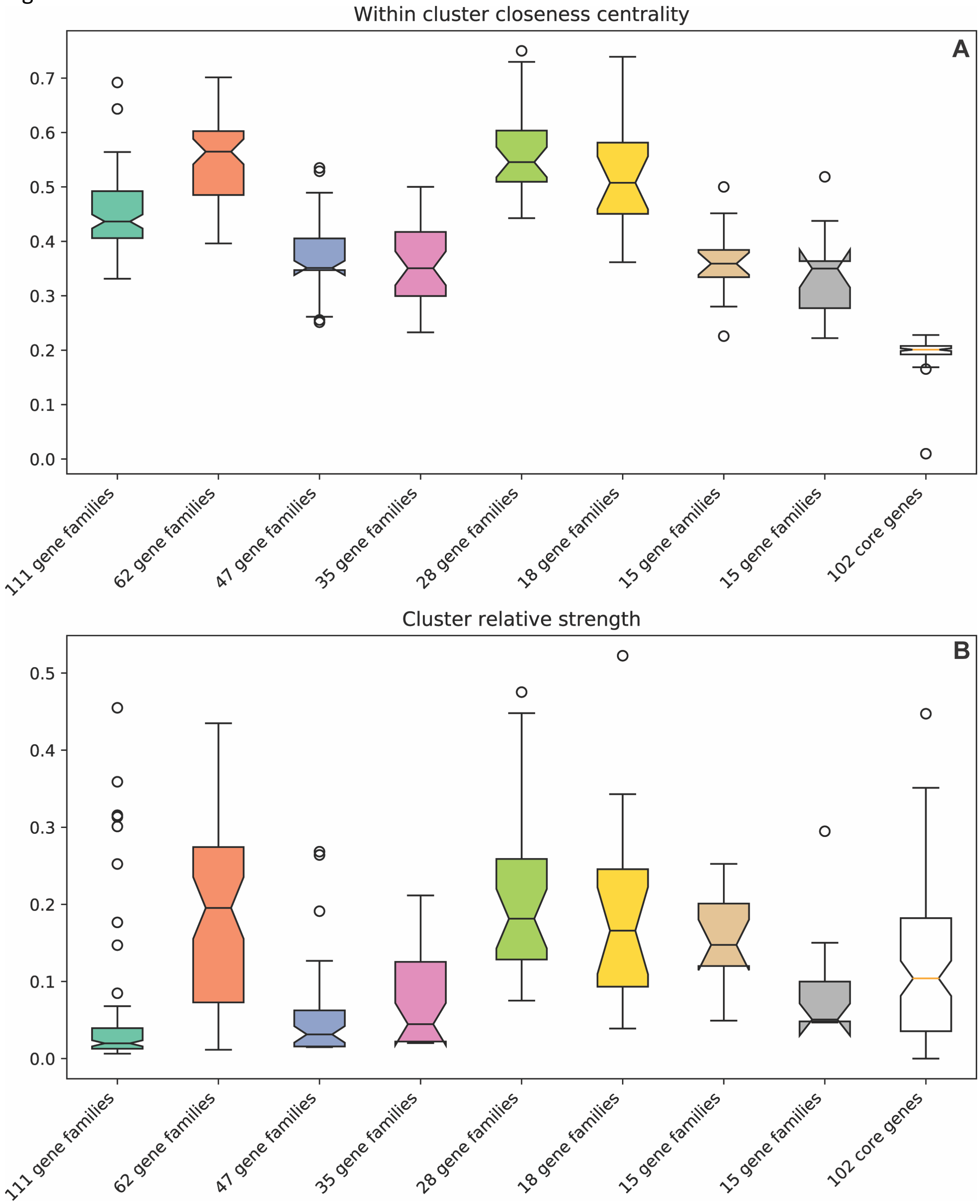

Representation of cluster robustness metrics. A) closeness centrality distributions calculated from subgraphs comprised by nodes from each co-evolving cluster independently. B) Weighted degree centrality, a.k.a. strength, divided by cluster size calculated from cluster specific subgraphs. In order, boxplots represent values estimated from cluster#3, cluster#4, cluster#2, cluster#0, cluster#5, cluster#8, cluster#1, and cluster#15.

Fig S4

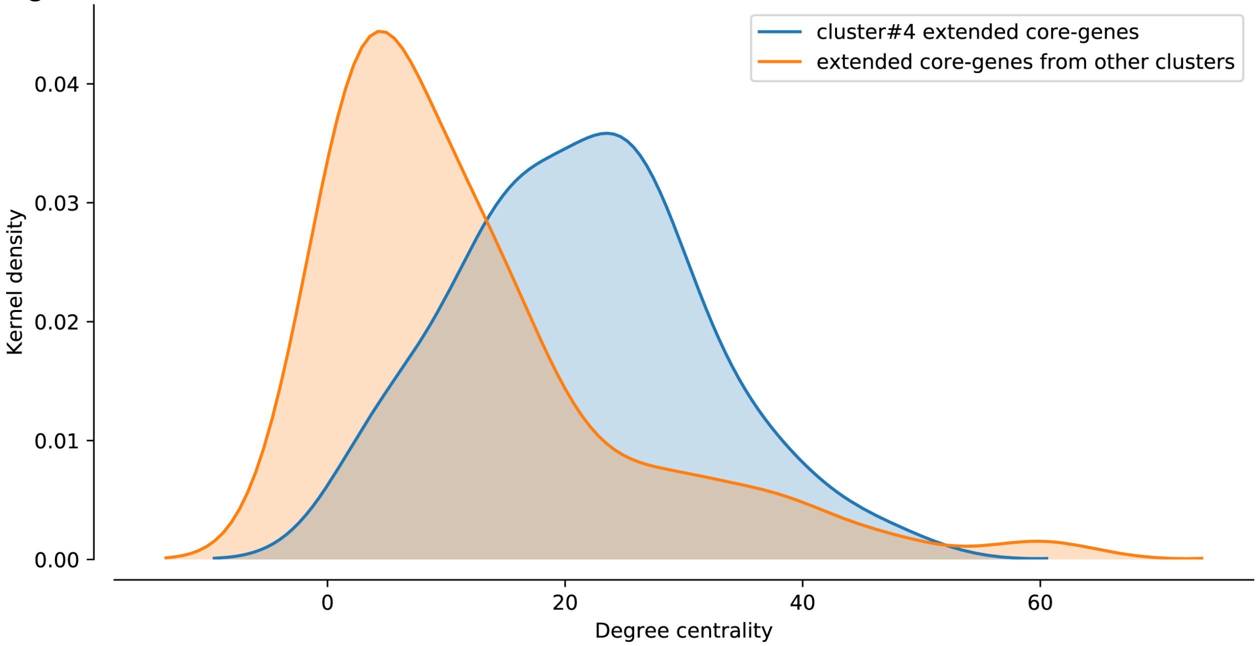

Degree centrality distributions from cluster#4's extended core genes with all 102 extended core genes in blue, and from non-cluster#4's extended core genes with all 102 extended core genes in orange. Evolutionary histories of extended core genes present within cluster#4 are similar to significantly more extended core genes.

Fig. S5

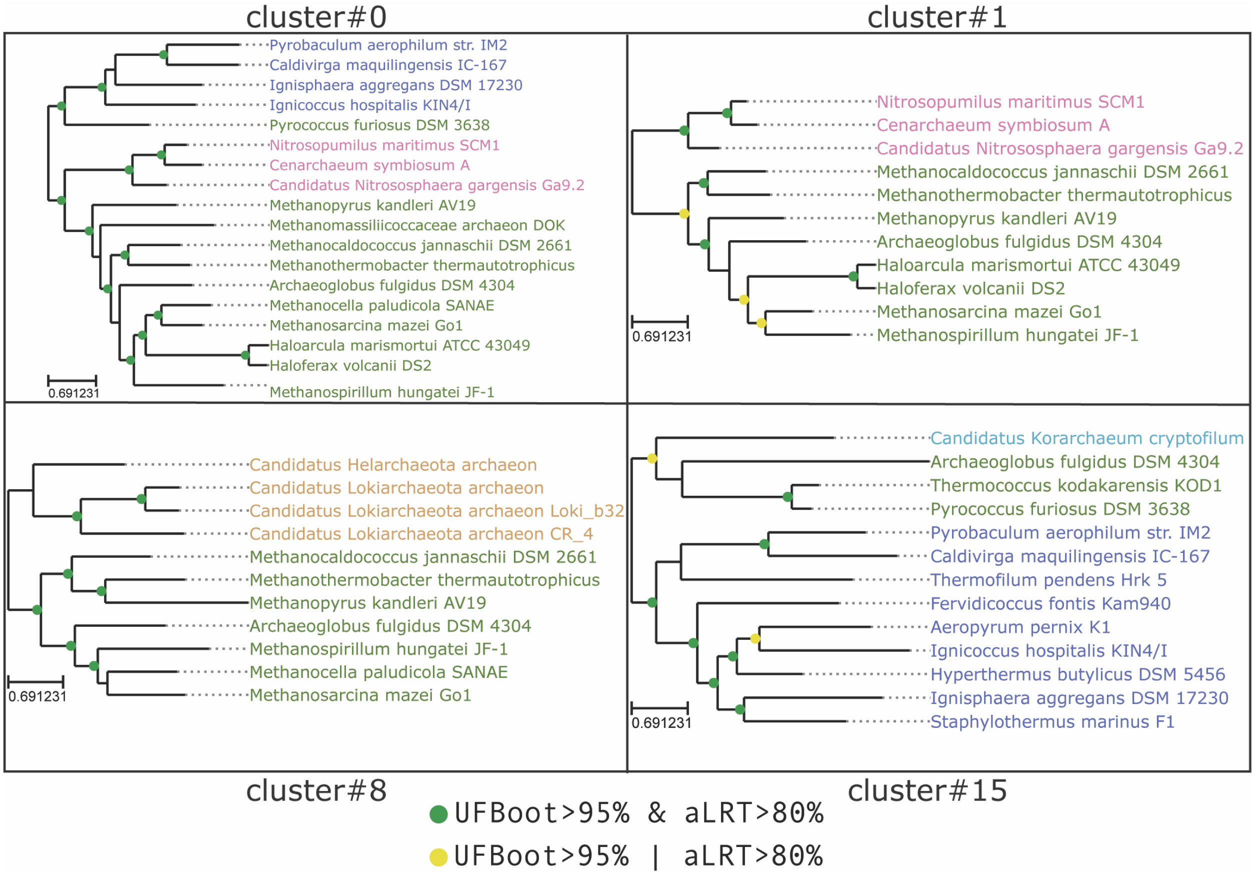

Phylogenetic trees reconstructed from concatenations of genes from each co-evolving cluster using LG+F+G+C60 evolutionary model. Genomes missing more than 25% of concatenated sites were removed before phylogenetic reconstruction. Leaf names are colored in the same structure as Fig. 7. Trees from each co-evolving cluster were independently rooted using MAD [1].

Fig. S6

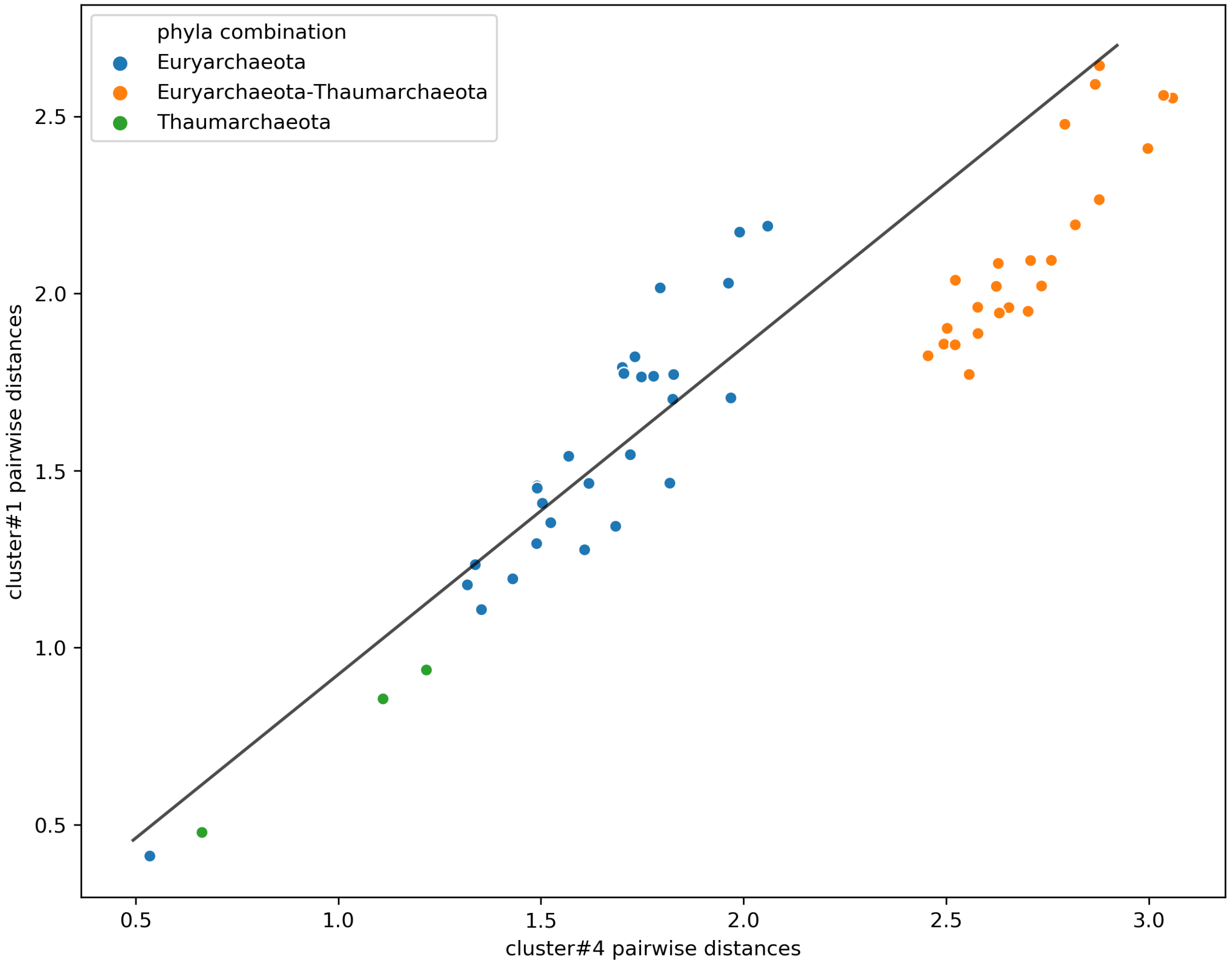

wODR between pairwise distances calculated using concatenated sequences from CES cluster#4 and CES cluster#1. Datapoints corresponding to intra and inter-phylum distances were assigned weights ten and one, respectively. This weighting scheme was used to anchor the regression model using intra-phylum pairwise distances, which better reflect the estimated vertical signal, and evaluate how inter-phyla distances fit the expected linear association.

Fig. S7

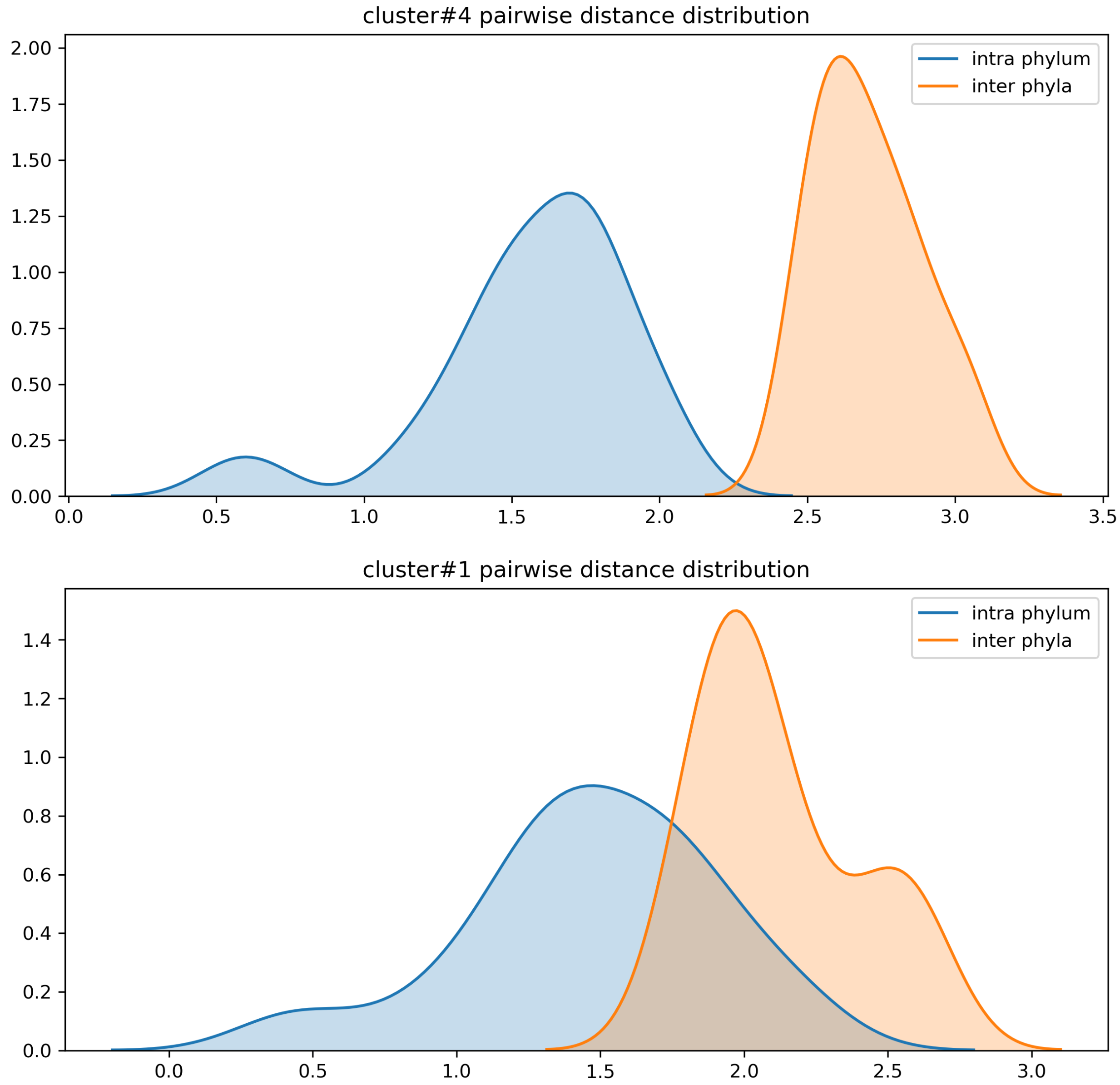

Distribution of intra-phylum distances in blue and inter-phyla distances in orange for cluster#4 and cluster#1, respectively. Pairwise distances were calculated using concatenated sequences from each CES cluster. Intra and inter-phyla distances are substantially different from each other in cluster#4 when compared to cluster#1. The smaller difference between intra and inter-phyla distances in cluster#1 is likely due a horizontal exchange of its co-evolving genes between both Euryarchaeota and Thaumarchaeota.

Fig. S8

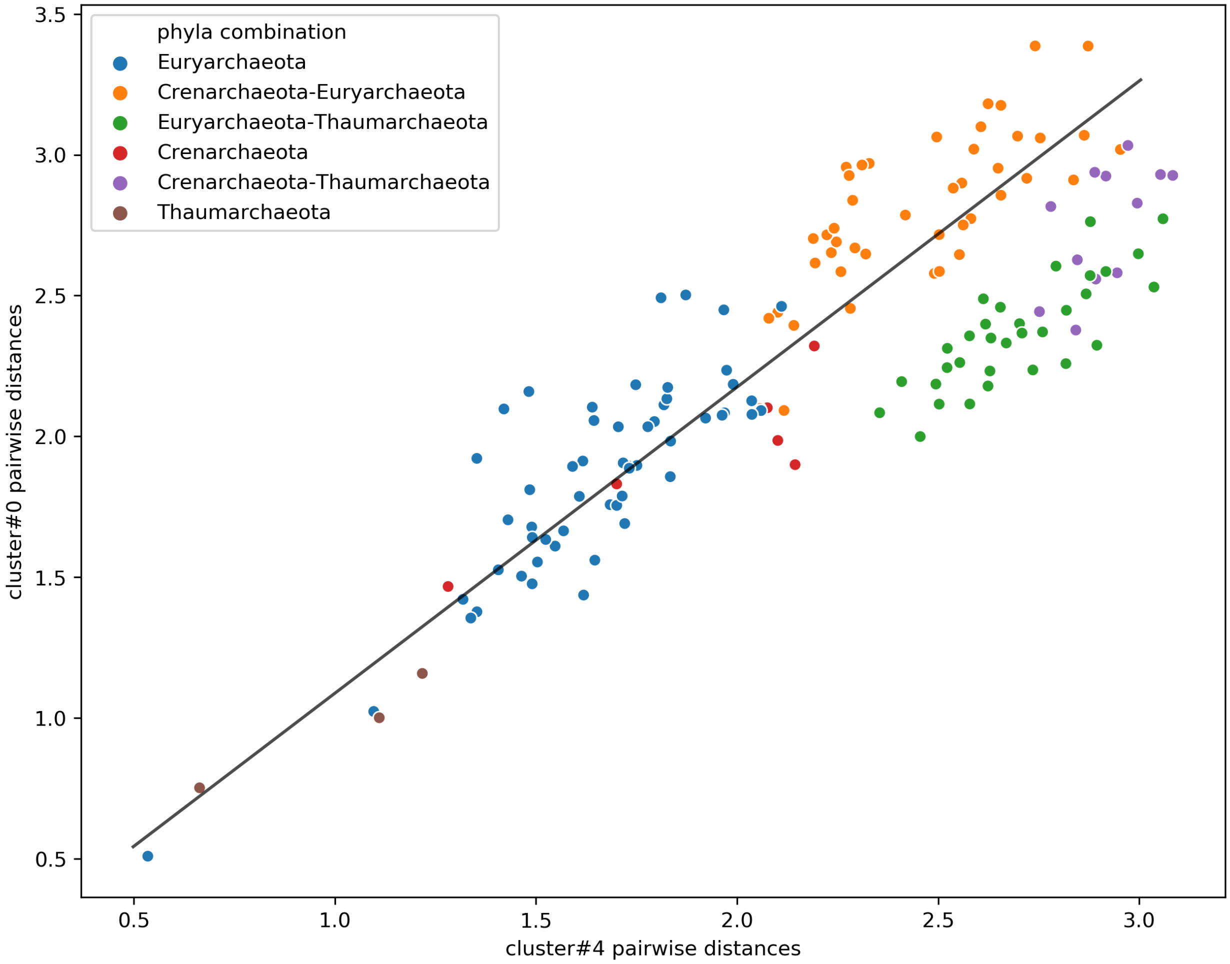

wODR between pairwise distances calculated from concatenated sequences from CES cluster#4 and CES cluster#0. Datapoints corresponding to intra and inter-phylum distances were assigned weights ten and one, respectively. This weighting scheme was used to anchor the regression model using intra-phylum pairwise distances, which better reflect the estimated vertical signal, and evaluate how inter-phyla distances fit the expected linear association.

Fig. S9

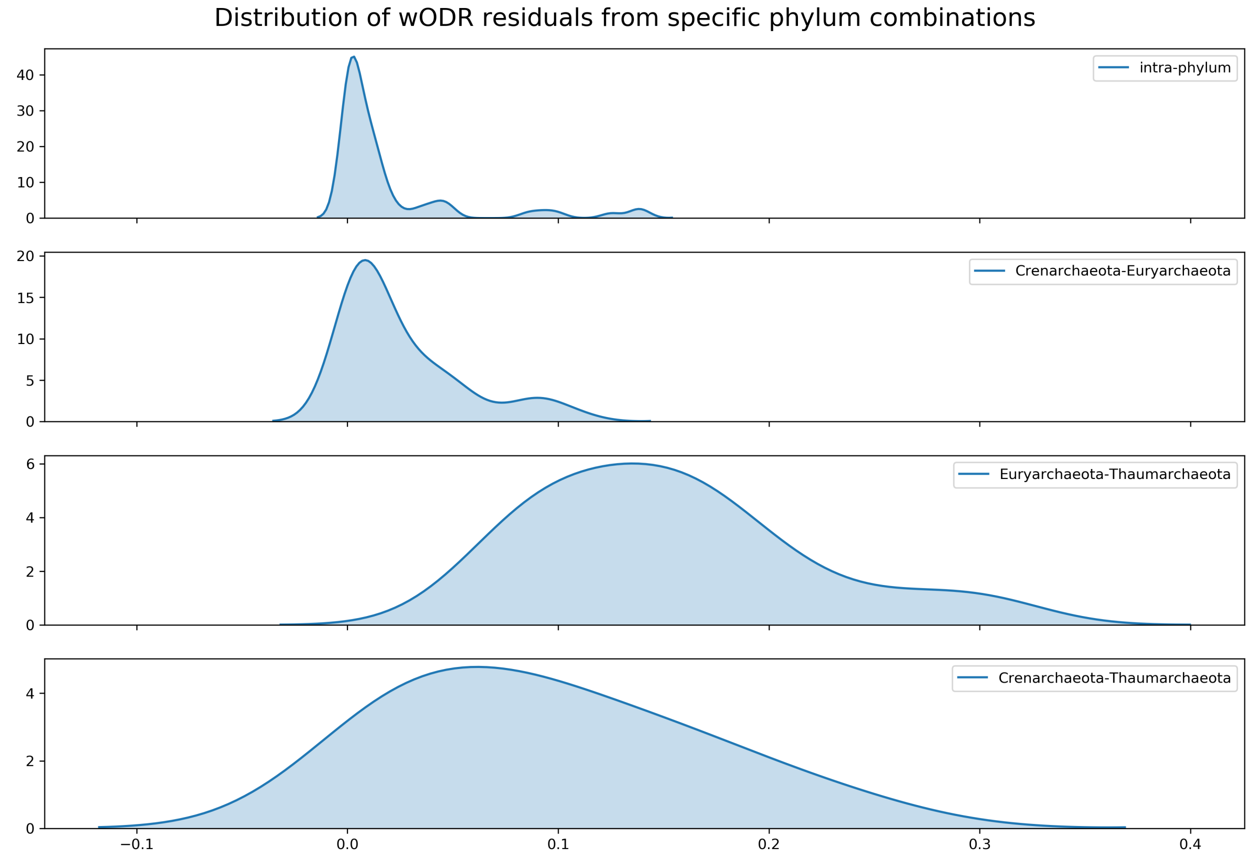

Distribution of wODR residuals calculated using concatenated sequences from CES cluster#4 and CES cluster#0 by intra-phylum distances and each combination of two phyla in CES cluster#0. Residuals were estimated from wODR fitted using intra and inter-phylum distances weights of ten and one, respectively. Using Common Language effect size statistics, Creanarchaeota-Euryarchaeota displayed very similar residuals ($f = 0.62$), while Euryarchaeota-Thaumarchaeota and Creanarchaeota-Thaumarchaeota displayed much larger residuals ($f = 0.97$ and $f = 0.90$, respectively).

Fig. S10

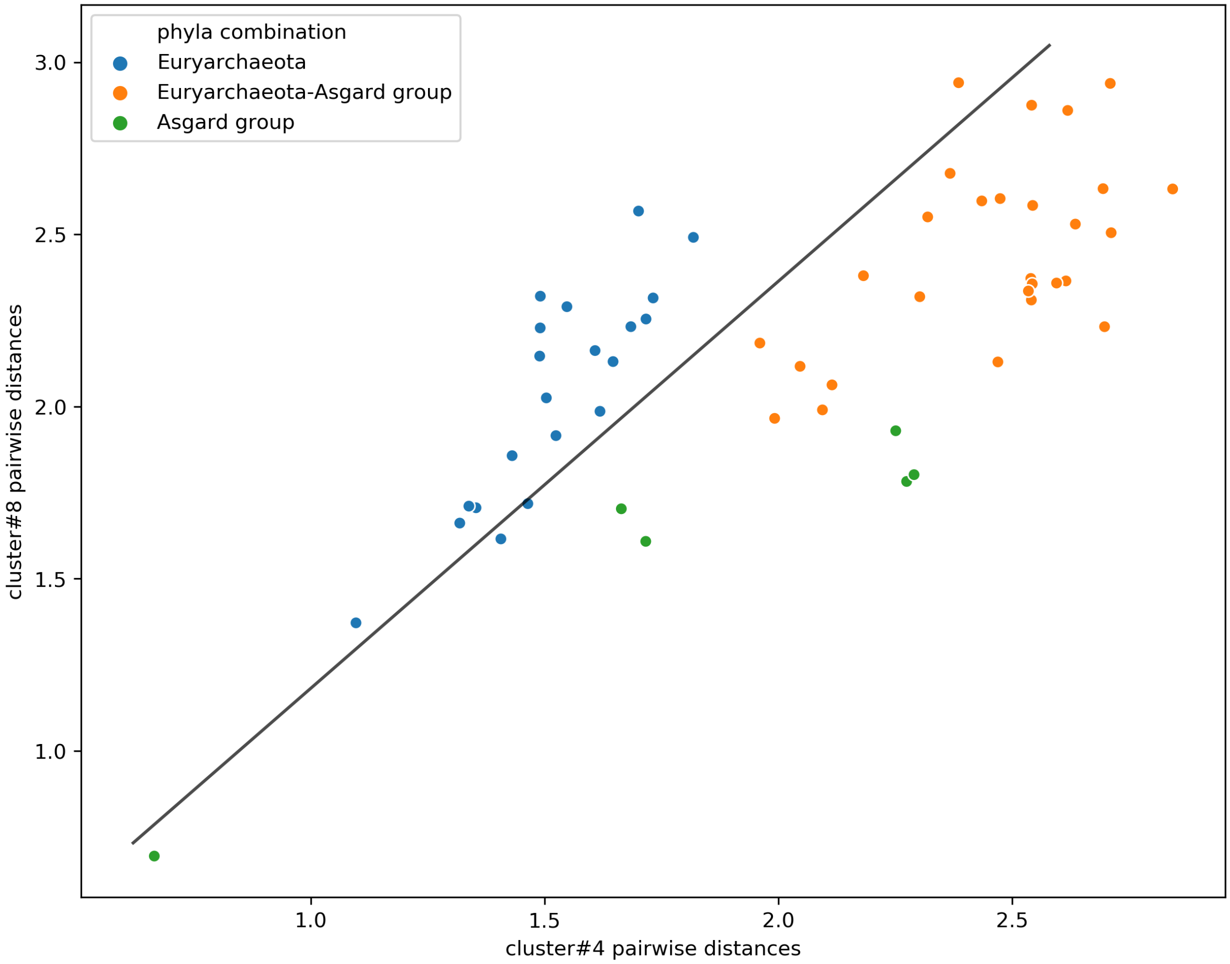

wODR between pairwise distances calculated using concatenated sequences from CES cluster#8 and CES cluster#0. Datapoints corresponding to intra and inter-phylum distances were assigned weights ten and one, respectively. This weighting scheme was used to anchor the regression model using intra-phylum pairwise distances, which better reflect the estimated vertical signal, and evaluate how inter-phyla distances fit the expected linear association.

Fig. S11

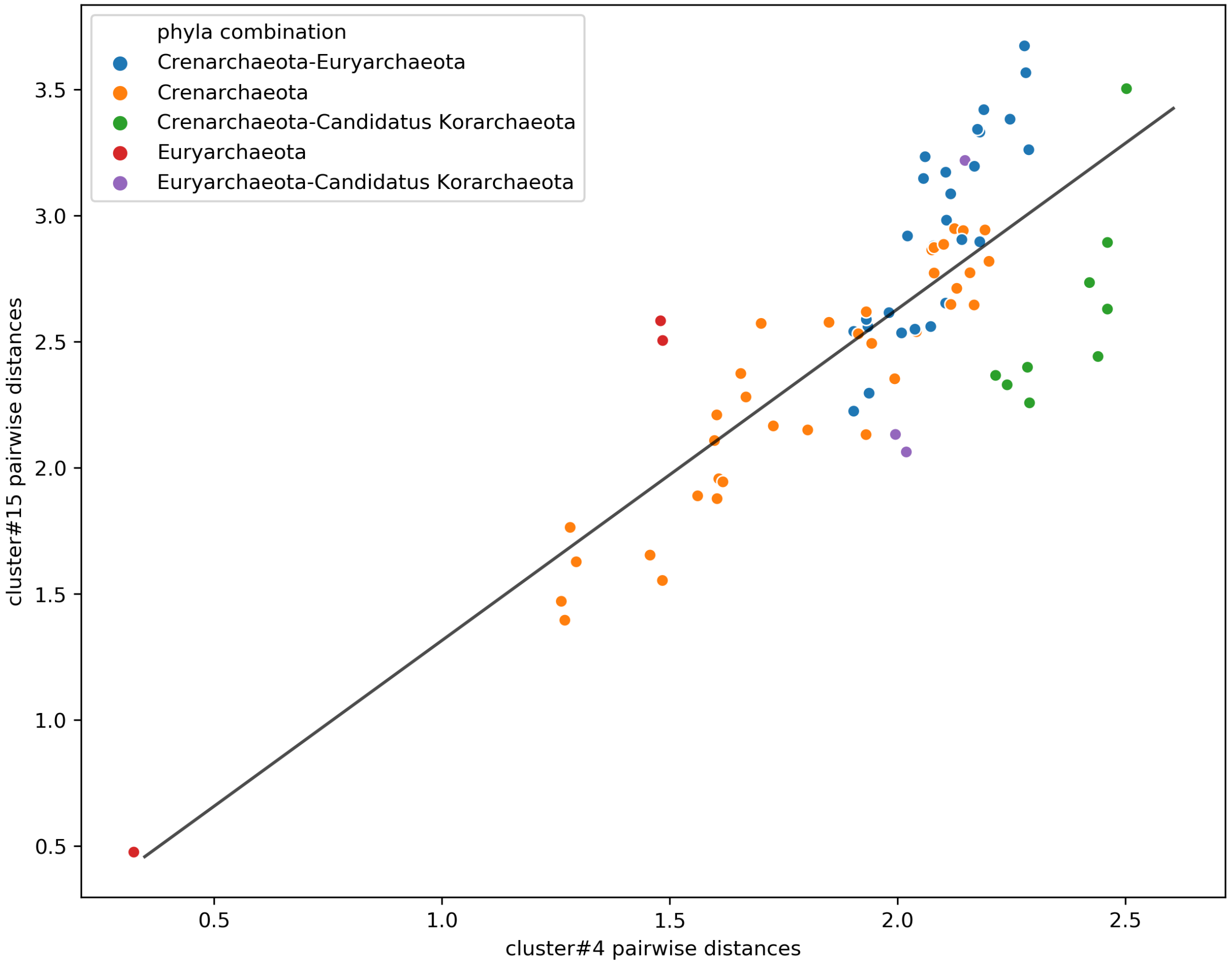

wODR between pairwise distances calculated using concatenated sequences from CES cluster#8 and CES cluster#0. Datapoints corresponding to intra and inter-phylum distances were assigned weights ten and one, respectively. This weighting scheme was used to anchor the regression model using intra-phylum pairwise distances, which better reflect the estimated vertical signal, and evaluate how inter-phyla distances fit the expected linear association.

Fig. S12

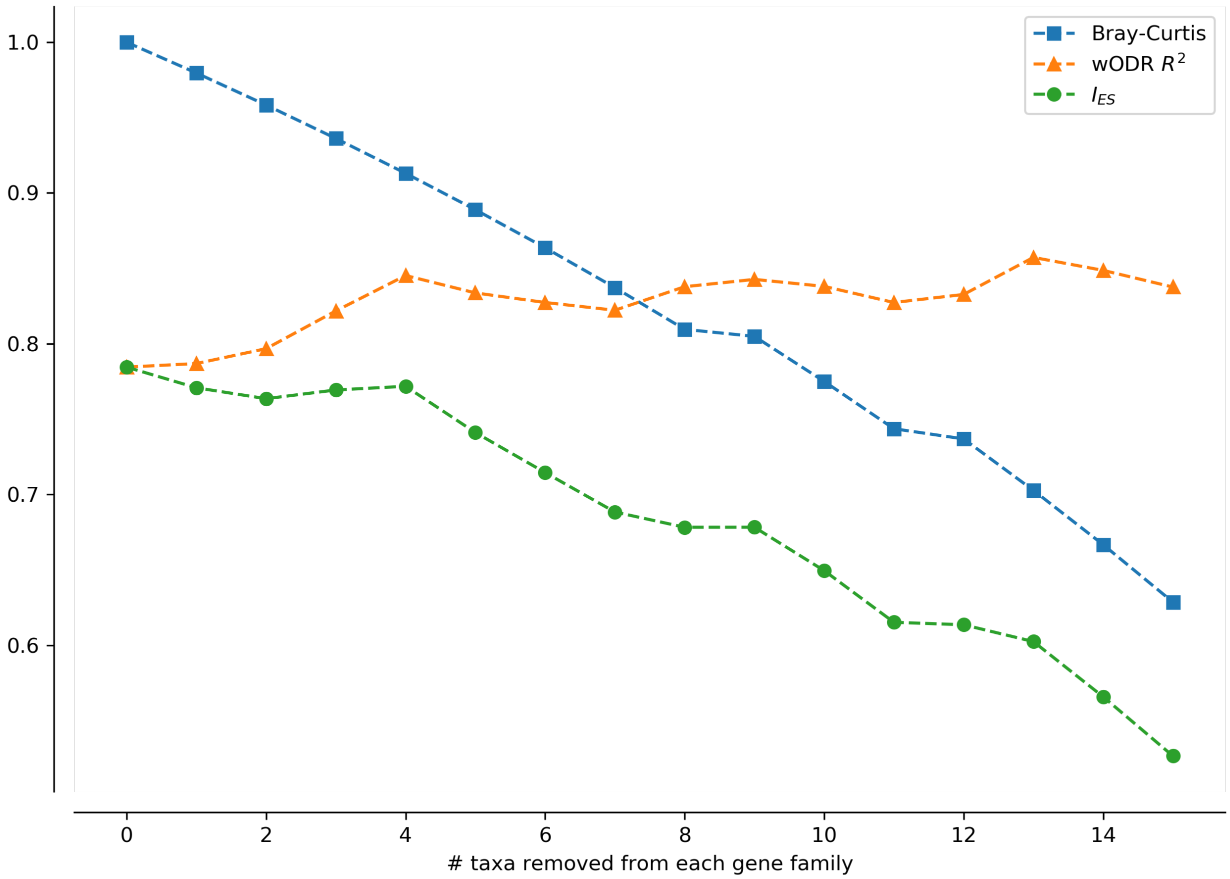

Stepwise changes to three metrics estimating similarities between evolutionary histories of gene families as sampled taxa randomly decrease. Bray-Curtis (blue) steadily decreases as consequence of dissimilar genome occurrences. wODR $\mathrm{R}^2$ (orange) tends to increase as taxa representing deviations between evolutionary histories are randomly removed. $I_{ES}$ (green) display an overall downward trend, although in a slower pace than Bray-Curtis, induced by differential gene losses between gene families. Both gene families contain 50 taxa, and their phylogenies diverge from each other by 5 SPR transformations.
